# Single-Molecule FRET Reveals Two T4 Phage MR Complex Exonuclease States Regulated by the Mre11 Dimer Interface

**DOI:** 10.1101/2025.07.29.667502

**Authors:** Jennifer B. Coats, Tibebe Teklemariam, Yang Gao, Scott W. Nelson

## Abstract

The Mre11/Rad50 (MR) complex serves as an initial responder to double-stranded DNA breaks and is conserved across all domains of life. Rad50 possesses ATPase activity that provides energy for the nuclease activities of Mre11. To elucidate the variation in exonuclease activity across the T4 phage MR complex population, we conducted smFRET studies utilizing a dual-labeled Cy3 and Cy5 double-stranded DNA substrate. Our findings revealed that the wild-type (WT) MR complex population was predominantly divided into two distinct rates of exonuclease activity. The majority of the population (94%) exhibited rapid exonuclease activity, with an average rate exceeding 20 nucleotides per second, while the remaining fraction showed substantially slower exonuclease activity, averaging 0.50 nucleotides per second. Furthermore, we compared the activity of the WT population to a mutant complex harboring the Mre11 mutation L101D, which has previously been shown to disrupt the Mre11 dimer interface. The MR complex with the L101D mutation exhibited a much higher percentage of the population displaying slow exonuclease activity (78%), while the remaining population exhibited fast exonuclease activity at a rate similar to that of the WT population. Additional activity parameters were evaluated, such as pause times and transition amplitudes, under various enzyme concentrations, revealing further differences between the WT and L101D MR complexes. Our results provide new insights into the molecular mechanisms of MR complex exonuclease activity and suggest that the complex may adopt different conformations with distinct kinetic properties.

## INTRODUCTION

The genomic stability of a cell is constantly challenged by several endogenous and exogenous factors including ionizing radiation, reactive oxygen species, and replicative stress that can lead to multiple forms of DNA damage^1^. The severity of the type of damage is often judged by the ability of a cell to repair the damage without inducing a mutation. Double-stranded DNA breaks (DSBs) are considered one of the most severe forms of damage because neither of the original strands remains intact to serve as a template for repair^2^. The loss of genomic stability through the failure to repair breaks in the DNA is a driving force behind many oncogenic events or can lead to cellular apoptosis. Cells use non-homologous end joining (NHEJ), microhomology-mediated end joining (MMEJ), and homologous recombination (HR) as the primary repair pathways for DSBs^2–4^. In all three of these pathways, the Mre11/Rad50 (MR) complex is a first responder that plays a role in signaling to other DNA repair enzymes, tethering of the loose DNA ends together, and resectioning through the nuclease activity of Mre11^5^.

Mre11 is a Mn^2+^ dependent nuclease that is part of the metallophosphatase superfamily. Within the MR complex, Mre11 forms a dimer with 3’ to 5’ exonuclease activity and ssDNA endonuclease activity that contributes to the formation of the 3’ overhang during resectioning^6,7^. Along with the nuclease active site, Mre11 has the capping domain which distinguishes between double-stranded and single-stranded DNA substrates. A helix-loop-helix motif at the C-terminal binds to Rad50 near the base of Rad50’s coiled-coil domain which extends away from the catalytic head of the tetrameric complex^8^. Rad50 dimerizes within the complex through the CXXC motif at the apex of the coiled-coil which forms a zinc tetrathiolate better known as the zinc hook; a characteristic which is unique within the ATP-binding cassette (ABC) ATPase superfamily^9^.

The MR complex is evolutionarily conserved across all domains of life. In eukaryotes, the MR complex forms a hexamer with two copies of Nbs1 (Xrs2 in yeast), which is absent in prokaryotes and archaea^6^. While the MR complex functions in DSB resectioning within eukaryotes, archaea, and bacteriophage systems, in prokaryotes this role is carried out by RecBCD^10,11^. The MR complex homolog, SbcCD, primarily cleaves hairpin loops within the genome^12,13^.

Bacteriophage T4 has proven to be a good model system for the study of genetic structure, genome replication and repair, and protein structure and function due to its minimalistic system requirements in comparison to higher organisms and its well understood biology^14,15^. The T4 MR complex (gp47/gp46) has the advantage of being easily purified without the need for truncation of Rad50’s coiled-coil region^16^. Compared to eukaryotes and prokaryotes, the T4 phage complex has a relatively short coiled-coil region which is thought to increase solubility and stability during purification. While the structure of the T4 phage MR complex has not been determined, sequence homology shows that T4 phage shares all of the basic structural features of eukaryotic and prokaryotic complexes including, in Rad50, the Walker A- and B-motifs, the D-loop, the H-loop, the Q-loop, and the CXXC motif at the apex of the coiled-coil and, in Mre11, the capping domain as well as five highly conserved phosphoesterase motifs which coordinate two Mn2+ ions at the active site^7,16,17^. Previous work has also shown that the T4 MR complex’s functionality in HR is comparable to eukaryotic complexes through Mre11’s single stranded endonuclease activity and 3’ to 5’ exonuclease activity as well as the increased activation of the complex’s ATPase activity by double-stranded DNA.

Previous ensemble kinetic studies have shown that the T4 MR complex in the presence of dsDNA will hydrolyze 3 to 4 ATP per second and excise ∼4 nucleotides per ATP^16^. Nuclease assays indicate that ATP is not necessary for the excision of the first nucleotide, suggesting that that ATP-hydrolysis primarily drives translocation along the DNA substrate. A previously studied T4 Mre11 mutant, L101D (Fig. 1), was used to elucidate the significance of the Mre11 dimer interface during ATPase and nuclease activity^18^. Mre11 dimerizes through the hydrophobic residues of a four-helix bundle which can be disrupted through the introduction of a charged residue such as L101D. In this study, binding affinity of L101D-Mre11 to Rad50 and ssDNA endonuclease activity of the L101D-MR complex were shown to be equivalent to the WT-MR construct. The presence of L101D had a significant impact on ATPase activity decreasing *k_c_*_at_ by 10-fold in comparison to WT. While the L101D-MR complex showed an increase in productive assembly to initiate nuclease activity, a decrease in the rate of exonuclease activity was observed. These results indicate that the L101D mutation has either reduced translocation or disrupted processivity.

**Figure 1.**
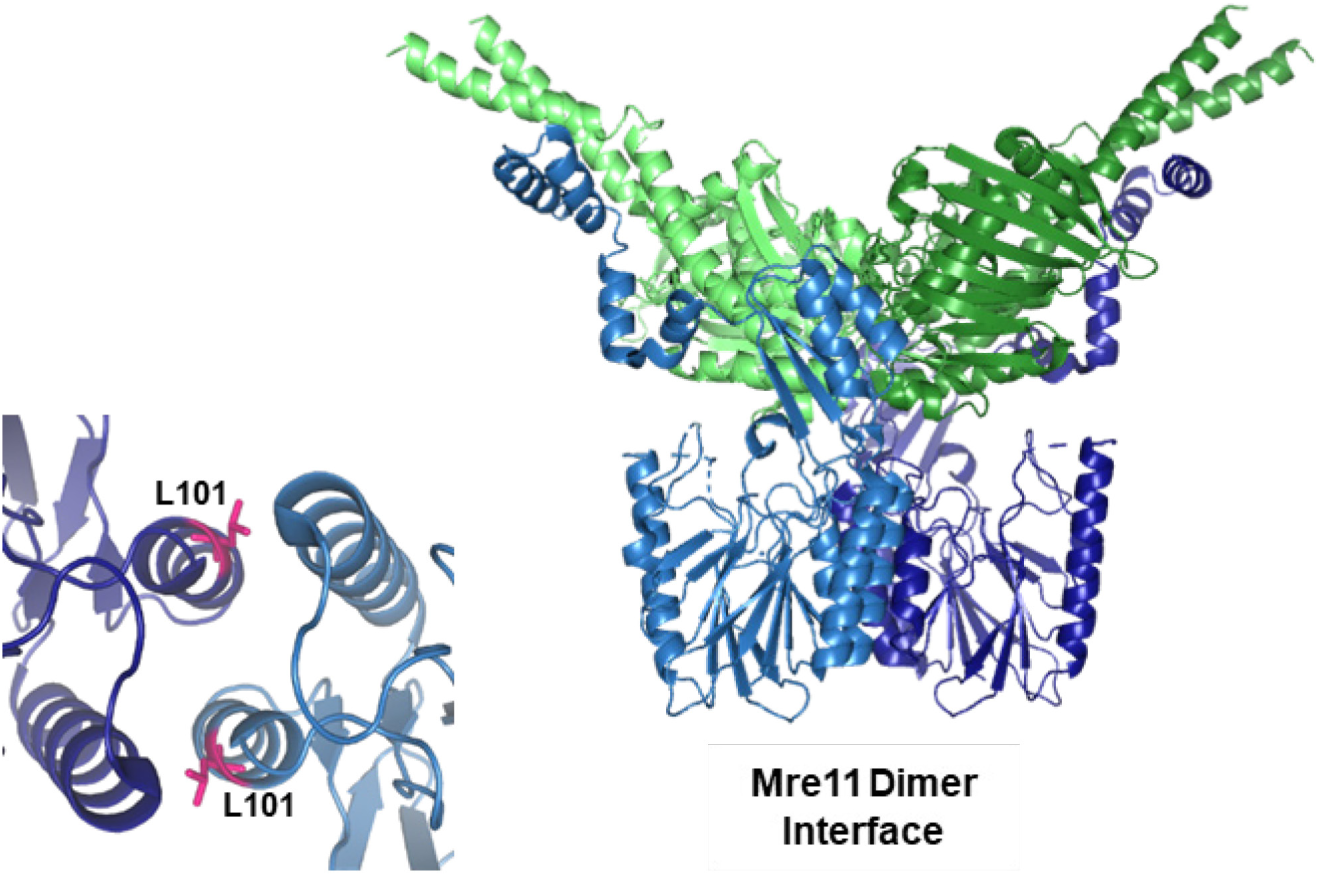
Example WT-MR complex structure from *T. maritima* showing the position of L101. The Rad50 subunits are colored green and the Mre11 subunits are colored blue with the interface indicated. An enhanced view of L101’s position at the dimer interface is shown at the bottom left. The crystal structure was retrieved from Protein Data Bank with code 3THO^25^. Figure adapted from Albrecht *et al.* 2012

Ensemble assays are limited in their scope to focus only on averages in kinetic rates across an enzyme population, which can limit our understanding to the phenomenological. To observe the heterogeneities of subpopulations with non-uniform kinetics, a single-molecule approach must be used. In this paper, we aim to better understand the variation in kinetics across the MR complex population by comparing the single-molecule kinetics of WT-MR to that of the mutant complex comprised of L101D-Mre11 and WT-Rad50. We analyze the population distribution of nuclease rates, processivity, and productive binding time to determine how the disengaged Mre11 dimer interface impacts different facets of the kinetic mechanism.

## MATERIALS AND METHODS

### Materials

Oligonucleotides were purchased from either Integrated DNA Technologies (IDT) or the Iowa State University DNA Facility. Labeled DNA substrates for FRET studies were purchased only from IDT. Glucose oxidase from *Aspergillus niger*, catalase from bovine liver, streptavidin from *Streptomyces avidinii*, and Trolox (6-hydroxy-2,5,7,8-tetramethylchroman-2-carboxylic acid) used in FRET studies were sourced from Sigma-Aldrich. The coupling enzymes pyruvate kinase and lactate dehydrogenase from rabbit muscle (PK/LDH) and nicotinamide adenine dinucleotide (NADH) used in ATPase assays also came from Sigma-Aldrich.

Phosphoenolpyruvate (PEP) was purchased from Alfa Aesar, ATP from Spectrum Chemical, (3-Aminopropyl)triethoxysilane (APTES) from Tokyo Chemical Industry, and mPEG-SVA MW 5000 (both biotinylated and unbiotinylated) from Laysan Bio. Antibiotics used in purification (Ampicillin and Kanamycin) were purchase from Gold Biotechnology. All other chemicals and reagents were sourced through Fisher Scientific.

### Purification of WT Rad50, WT Mre11, and L101D Mre11

Purification of Rad50 and Mre11, both WT and mutants, was completed as described previously by Herdendorf *et al.* (2010 and 2011)^16,17^. T4 Rad50 in a pET28b vector was transformed into BL21 (DE3) cells. From overnight starter cultures, 1-liter cultures were inoculated and grown at 37°C to an OD of 0.6 at 600 nm. The cultures were then induced with 0.3 mM IPTG at 16°C overnight. Cells were harvested by centrifugation and resuspended in a buffer composed of 500 mM NaCl, 40 mM Tris, and 10 % glycerol at pH 7.8. After lysis using homogenization and centrifugation to separate the supernatant from the pellet, a nickel column was used to purify the protein. The nickel column was washed with the same resuspension buffer followed by the resuspension buffer with 20 mM imidazole and then an overnight wash of the resuspension buffer with 1 M NaCl. The protein was then eluted with 150 mM imidazole. Dialysis was performed to reduce the imidazole concentration of the sample before aliquoting and flash freezing for storage at -80°C.

T4 Mre11 in pTYB1 was transformed into BL21 (DE3) cells. Cell cultures were grown and lysed similarly to the description above. Cells were resuspended in a chitin binding buffer composed of 1 mM EDTA, 500 mM NaCl, and 20 mM Tris at pH 7.8. The supernatant after lysis and centrifugation was applied to a chitin column and, after washing with binding buffer the protein was eluted with 75 mM beta mercaptoethanol, 200 mM NaCl, and 20 mM Tris at pH 7.8. Dialysis was completed to remove the beta mercaptoethanol and then aliquots were frozen at - 80°C.

### Design of the Substrate DNA

The dsDNA substrate was adapted from Lee *et al.*^19^ with minor modifications to the design and sequence. The substrate was 50bp long and attached to the PEGylated surface of the slide through the biotinylated 5’ end of the unlabeled strand. A thiophosphate linkage was placed 12 bases away from the 5’ end of the unlabeled strand to stop exonuclease activity and avoid dissociation of the labeled strand from the slide surface. On the labeled strand, the Cy3 was positioned at the 5’ end and Cy5 was positioned 24 bases away.

### Oligonucleotide Sequences

Labeled strand: 5’-/5Cy3/ GAA CTT AAT AAT CTG TCT CTC CAG / iCy5/ GAA CCA ATC GCG GAC GAG CAC ACC AG-3’

Unlabeled strand: 5’-/Biosg/ CTG GTG TGC TCG* TCC GCG ATT GGT TCC TGG AGA GAC AGA TTA TTA AGT TC-3’

Unlabeled product mimic strand: 5’-/Biosg/ CTG GTG TGC TCG TCC GCG ATT GGT-3’

### Preparation of the Slides for FRET

Procedures for cleaning the quartz slides used in imaging were adapted from Chandradoss *et al.*^20^ with minor modifications. Slides being reused from previous imaging sessions would be stored in acetone until ready to be cleaned to mitigate the removal of epoxy and other residue still on the slides. To initiate the cleaning procedure, 8 to 12 slides were boiled in 400 mL pyranic acid (300 mL sulfuric acid and 100 mL hydrogen peroxide) for 25 minutes. After rinsing thoroughly with Milli-Q water, the slides were placed with the coverslips in a 1 M KOH solution and sonicated for 45 minutes to etch the quartz surface. After rinsing in water, the slides were once again placed in a freshly made solution of pyranic acid (without heating) for 25 minutes. Coverslips were never exposed to pyranic acid during the cleaning procedure. After rinsing thoroughly with Milli-Q water, the slides were once again combined with the coverslips to be rinsed with methanol four times. The slides and coverslips were sonicated in the methanol for 15 minutes during the third rinse. After removal of the final wash with methanol, the slides and coverslips were prepared for PEGylation with a silanization solution comprising of 2.5 % glacial acetic acid and 0.5 % APTES (3-aminopropyl trimethoxysilane) suspended in methanol. The slides and coverslips would sit in this solution for 10 minutes, be sonicated for 1 minute, and then left to sit an additional 15 minutes before removing the silanization solution and rinsing, first with methanol and finally with Milli-Q water. Each slide would then be PEGylated with 0.2 mg of biotinylated NHS-ester PEG 5000 and 8 mg of non-biotinylated NHS-ester PEG 5000 dissolved in 640 µL 0.1 M filtered NaHCO3 (pH 8.5). After pipetting this solution onto the surface of the slide, the coverslip is placed over it and the assembly is left overnight in the dark. The next day, the slides and coverslips were washed with Milli-Q water and dried with nitrogen gas. Slides were stored under vacuum in 50 mL centrifuge tubes until ready to image.

Another round of PEGylation would be completed the day of imaging. This was followed by preparation of the individual channels. Prior to cleaning, quartz slides were prepared for imaging by drilling ¾ mm holes along the two long edges of the slide. These holes were spaced 1.5-2 mm apart along the edge and paired with a counterpart symmetrically placed across from them creating an inlet and outlet pairing. After PEGylation the day of imaging, 1 mm thick strips of Scotch permanent double-sided sticky tape were placed between each pair of holes and along the outer edges of the first and last pairs before the coverslip was placed over the tape. This would create channels with one inlet and one outlet hole apiece. A small amount of epoxy was run along the length of the coverslip edge to complete enclosing the channels with the exception of the inlet and outlet. Tubing, approximately 4 cm in length, was placed in both the inlet and outlet holes and epoxy was used to seal the tubing in place.

Once the channels were fully assembled, 50 μL 0.05-0.1 mg/mL streptavidin was flowed through each channel. After a 5-minute incubation, 500 μL of wash buffer (50 mM Tris-HCl, 50 mM KCl, pH 7.8) was used to remove excess streptavidin. 200 μL of 40 pM biotinylated DNA substrate is flowed through each channel and left to incubate for 2 minutes before being washed away using the same wash buffer. The slide was covered to reduce light exposure until ready for imaging.

### Data Collection and Analysis

Data films were collected using an Andor iXon Ultra 888 EMCCD and Single v1.0.4. Two-minute videos were filmed with an exposure time of 0.4 seconds per frame, a signal gain of 500 and a data scaler of 150. Prior to flow of the enzyme and substrates into the channel, imaging buffer composed of 50 mM Tris-HCl (pH 7.8), 50 mM KCl, 10 mM MgCl_2_, 0.1 mg/ml BSA, 0.4 % glucose, 2 mM Trolox, 0.6 mg/ml glucose oxidase, and 0.07 mg/ml catalase is flowed into the chamber and a video confirming the initial state of the labeled DNA substrate is taken. Glucose oxidase and catalase are added fresh to the imaging buffer immediately before use. Videos of the exonuclease reactions began simultaneously with the flow of 140 μL of the imaging buffer with 400nM of the MR complex, 0.3 mM MnCl_2_, and 1 mM ATP into the channel at a rate of 280 μL/second (Fig. 2).

Videos were analyzed using iSMS: single-molecule FRET microscopy software to extract the acceptor/donor traces from the individual FRET-pairs and calculate the FRET trace^21,22^. After manually removing poor-quality spots based on brightness, photobleaching, and proximity to neighboring spots, traces were exported to be analyzed using BEAST (Bayesian Estimator of Abrupt change, Seasonality, and Trend), which was used for change-point detection and identification of regions with positive slopes indicating transitions^23^. The BEAST output was then manually compared with the acceptor and donor traces to confirm actual transitions as opposed to blinking or photobleaching. Because the time scale over which transitions occurred varied significantly within our data set, two approaches were used to capture all transitions. First, we identified regions with a greater than 15 percent probability of having a positive slope over multiple frames, which allows identification of transitions that occur over a longer timescale. Second, we identified individual frames where the slope varied by more than one standard deviation away from the average slope of all frames in the movie. This effectively captures faster transitions that occur in five frames or less and are not recognized by the first method for identifying transitions.

Transitions were defined by their beginning and ending timepoints as well as their initial and final FRET. While product mimic data indicated a very slight sigmoidal curve for the conversion between nucleotides excised and FRET (Supplemental Fig. S1), our data falls primarily in the linear region. Because of this potential sigmoidicity, we limited our data to 18 nucleotides excised rather than 24 as our substrate would allow. All transitions greater than 18 nucleotides excised are grouped together and calculations performed are treated as limits rather than actual representative values.

## RESULTS

### Control Experiments

Prior to starting smFRET experiments, the impact of DNA labeled with fluorescent dyes on MR complex activity was tested by comparing the dual-labeled substrate to a 2-aminopurine labeled DNA substrate that has been used previously^16,18,24^. Results from ensemble assays showed that the fluorescent labels did not hinder activity and improved the sensitivity of the assay compared to the 2-aminopurine construct. Whereas 2-aminopurine only reports final product formation, the FRET assay readily captures exonuclease reaction intermediates (Supplemental Fig. S2).

For the main analysis, which used 400 nM MR complex, 27 L101D-MR videos and 29 WT-MR videos were recorded, yielding 2253 L101D-MR exonuclease events and 915 WT-MR events. Since the amount of DNA substrate is the same for both conditions and the DNA affinity for both proteins are similar, the higher number of observed exonuclease events for L101D-MR likely reflects less nonproductive binding than WT-MR^18^. No exonuclease activity was observed when ATP or Mn^2+^ is omitted from the reaction.

A major limitation affecting the achievable frame rate and duration of smFRET data acquisition is dye photobleaching. Eight representative channels (four each for L101D-MR and WT-MR) were used to quantify and assess the impact that photobleaching might have on our ability to capture transitions over the duration of data collection. Approximately 20 percent of the traces had photobleaching by the end of the two-minute collection time frame and 10 percent had photobleaching events occur within the first 47 seconds (Supplemental Fig. S3). In this same 47 second time frame, 84 percent of transitions had been initiated and 43 percent had been completed. As expected, the time distribution of photobleaching was independent of the mutant being tested.

### Exonuclease Events Fall Primarily into Two Distinct Subpopulations Based on Rate

Previous results using short DNA substrates (50 bp or less) indicate that the exonuclease reaction of the T4-MR complex is composed of a rate-limiting productive assembly step followed by exonuclease, ATP hydrolysis, and ssDNA translocation steps^16,18^. Within a FRET time trace, binding and productive assembly to initiate exonuclease activity is measured as the initial and subsequent dwell times where no change in FRET is observed prior to exonuclease activity. Exonuclease events are observed as transitions in FRET during a trace in which the increase in amplitude of the FRET change correlates to the number of nucleotides excised. The exonuclease rate is calculated as the slope of the FRET trace during a transition converted to nucleotides per second. ^25^

While most observed WT-MR exonuclease events have durations of less than one second, a subset proceed at a slower rate over several seconds, with a few extending beyond one minute. (Fig. 2 and 3). When compared to the L101D-MR mutant complex with a disengaged dimer interface, we similarly observed both fast and slow exonuclease subpopulations. Negative-stain transmission EM images (Supplemental Fig. S4) of the WT T4-MR complex with ATPγS in the absence of DNA reveal that a significant fraction of complexes adopt an open Mre11 dimer interface prior to DNA binding, suggesting that some may fail to form or maintain dimerization during activity.

**Figure 2.**
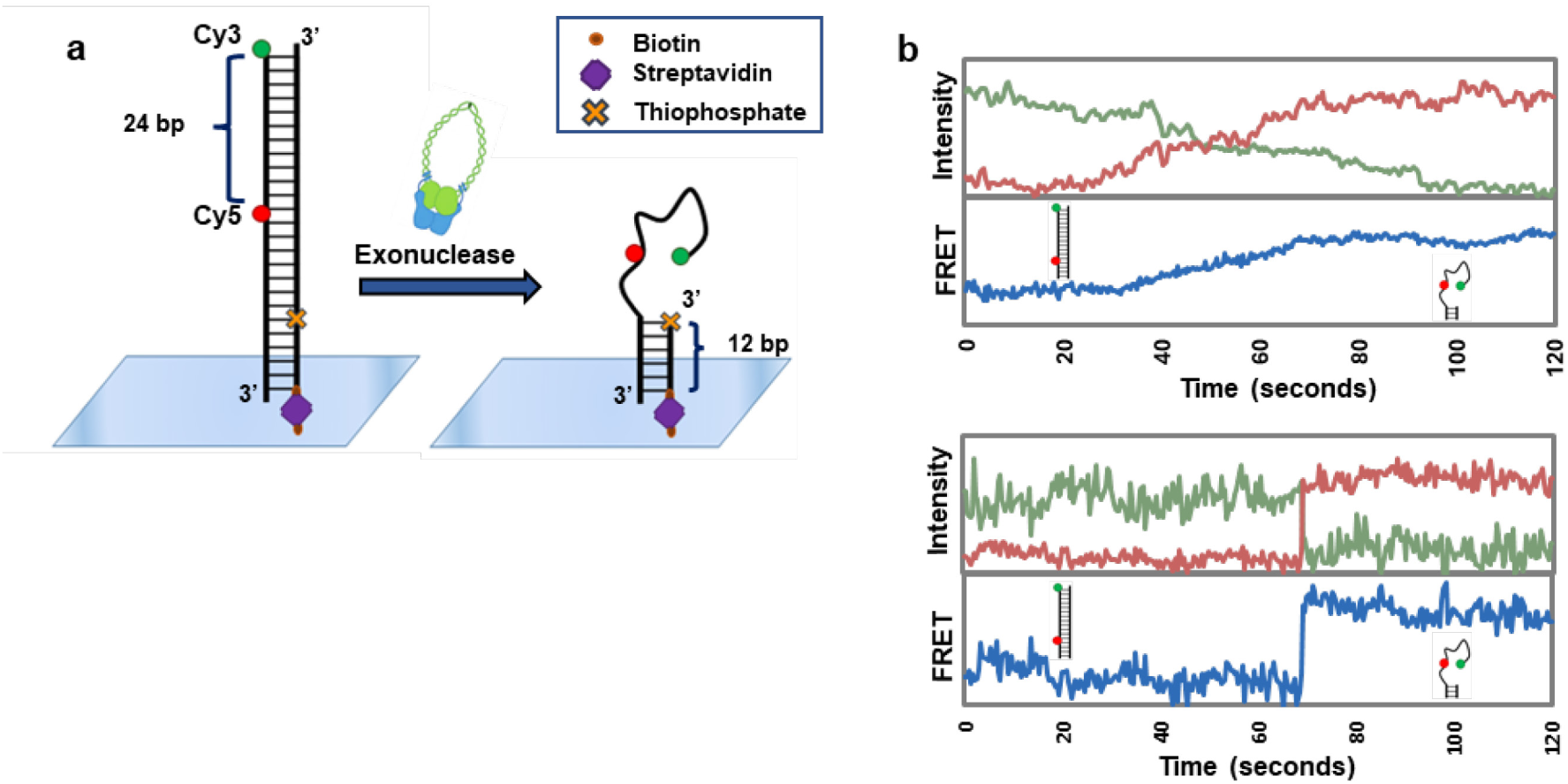
Experimental setup and trace variations a) Schematic of experimental setup on the slide before and after exonuclease activity. b) Example traces of the slow (top) and fast (bottom) exonuclease rate subpopulations observed. The change in fluorescence for Cy5 (red) and Cy3 (green) are shown normalized to 1. Change in FRET (blue) over time is also on a scale of 0 to 1.

Additional smFRET experiments were performed using a truncated Mre11 mutant missing the Rad50 binding domain and linker (ΔC289) in the absence of Rad50 (Supplemental Fig. S5). Previous work has shown that (ΔC289) has increased binding affinity to DNA compared to WT-Mre11 and, in bulk assays, has exonuclease activity that is detectable at much lower protein concentrations than WT-Mre11 alone^26^. Without Rad50, T4 phage Mre11 is monomeric and has negligible binding affinity for DNA^18,26^. The binding affinity for DNA of the complex comes primarily from Rad50’s head domain^18,27^. The smFRET exonuclease reactions with (ΔC289) indicate a slow exonuclease activity very similar to that of the slow subpopulation found with WT-MR. This suggests that the disengaged dimer interface of Mre11 in the presence of Rad50 is likely the result of Mre11 working as a monomer with increased binding affinity due to the presence of Rad50.

With L101D-MR, the population trend shifted toward the slower exonuclease activity compared to WT-MR. To categorize exonuclease rates into distinct populations, a cutoff of 1.35 nucleotides per second was selected based on a presumed Gaussian distribution of the rate data on a logarithmic scale (Fig. 3a). Using this criterion, 94 % of WT-MR events were classified as fast transitions, compared to only 22% of L101D-MR events (Fig. 3b). We also observed that the average exonuclease rates within the fast and slow transition subpopulations were increased for WT-MR (20.2 and 0.50 nucleotides/second, respectively) compared to L101D-MR (11.8 and 0.36 nucleotides/second, respectively) (Fig. 3c). The different rates likely indicate that, while L101D-MR can be used as an indicator of how a disengaged dimer interface impacts activity within WT-MR, the conformation of the disengaged dimer interface may differ to some extent from that of the WT-MR slow rate subpopulation.

**Figure 3.**
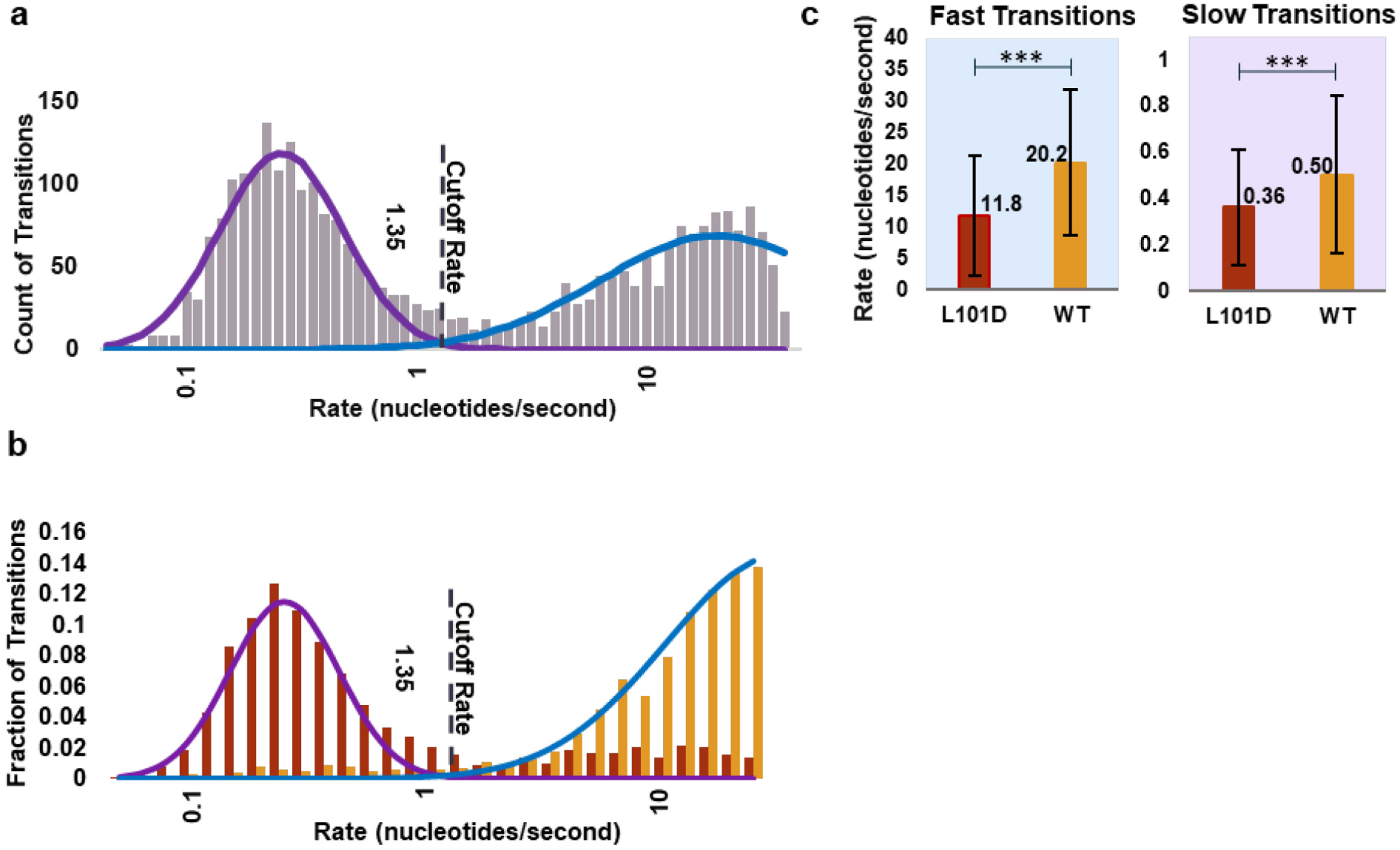
Rate population cutoff determination a) Data fitted to find cutoff between exonuclease rate subpopulations. All data fitted to two curves assuming a Gaussian distribution. A cutoff rate of 1.35 nucleotides/second was calculated as the intersection of the two curves. b) The fit of the two major peaks from the L101D (red) and WT (orange) data fitted separately. Each was fitted to two curves (minor fitted curves not shown). The calculated curves cross at 1.64 nucleotides/second. c) Average rate of WT-MR and L101D-MR complexes within each rate population. Slow transitions (purple) are the subpopulation below 1.35 nucleotides per second and fast transitions (blue) are the subpopulation above 1.35 nucleotides per second. Error bars show distribution of one standard deviation. Statistical analysis of figure c described in Supplemental Table S1.

### Determination of Processivity Through Transition Amplitudes

The linear correlation between FRET intensity and nucleotides excised extends to 18 bases removed, limiting our ability to accurately determine exonuclease processivity beyond this range. For this reason, transitions corresponding to more than 18 nucleotides excised were represented as 18 nucleotides. Although these transitions accounted for only a small proportion of the dataset (5.8%) and fell within a narrow range (18–24 nucleotides), the reported processivity values should be interpreted as lower estimates rather than absolute values.

Pre-steady state kinetics indicate that the L101D-induced disengaged dimer interface causes either a reduced rate in translocation or limited processivity^18^. The smFRET data presented here suggests exonuclease events occurring at a fast rate (>1.35 nucleotides per second) are processive for both L101D-MR and WT-MR. If intermediate steps are involved, then they occur on a time scale shorter than 100 ms (Supplemental Fig. S6). The data also indicate that the slow subpopulation exonuclease events for both L101D-MR and WT-MR are processive and arise from reduced translocation, as reflected by a gradual, continuous fluorescence trajectory lacking the discrete transitions characteristic of stairstep patterns as seen in the example slow traces of Figure 2b.

In categorizing the exonuclease events by mutant and rate of activity, an analysis of variance (ANOVA)^28^ indicated that the effect of construct, L101D versus WT (p<.001), and rate, fast versus slow ( p<.001), had a significant impact on the number of nucleotides excised both when considered as independent factors and in combination (p<.001) (Supplemental Table 2). When further Tukey post-hoc analysis^29^ was completed, we found that WT-MR had no significant correlation (post hoc p>0.05) between processivity and exonuclease rate (9.8 nucleotides excised for fast transitions and 10.5 nucleotides excised for slow transitions) (Fig. 4, Supplemental Table 3 for post hoc analysis). L101D-MR, however, did show a correlation (post hoc p<.0001) between the two rate subpopulations and their average processivity with faster transitions having lower amplitudes (7.3 nucleotides excised) and slower transitions having higher amplitudes (12.6 nucleotides excised).

### Characterization of MR Complex Interactions with Blunt and Recessed DNA Ends Through Analysis of Initial and Subsequent Dwell Times

For statistical analysis, an ANOVA^28^ was run on the logarithmized (due to skew from normal) data displayed in Figure 5, taking into consideration the effects of construct (L101D-or WT-MR), step (initial or subsequent), rate subpopulation (fast or slow), and the interactions between these effects using R-Studio (Supplemental Table 4)^30^. All individual effects and pairwise interactions reached significance (p < 0.05), except for the three-way interaction. Tukey’s post hoc analysis was run following the ANOVA analysis for further insight (Supplemental Table 5)^29^.

**Figure 4.**
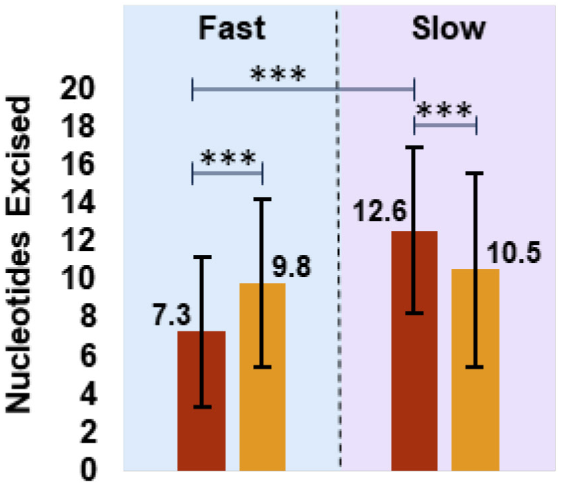
Average amplitudes based on construct and rate subpopulation for WT-MR (orange) and L101D-MR (red). Average number of nucleotides excised (amplitude) of transitions for WT-MR and L101D-MR divided into both fast and slow subpopulations. Amplitude changes notably for L101D-MR depending on the rate subpopulation but stays stagnant for WT-MR.

**Figure 5.**
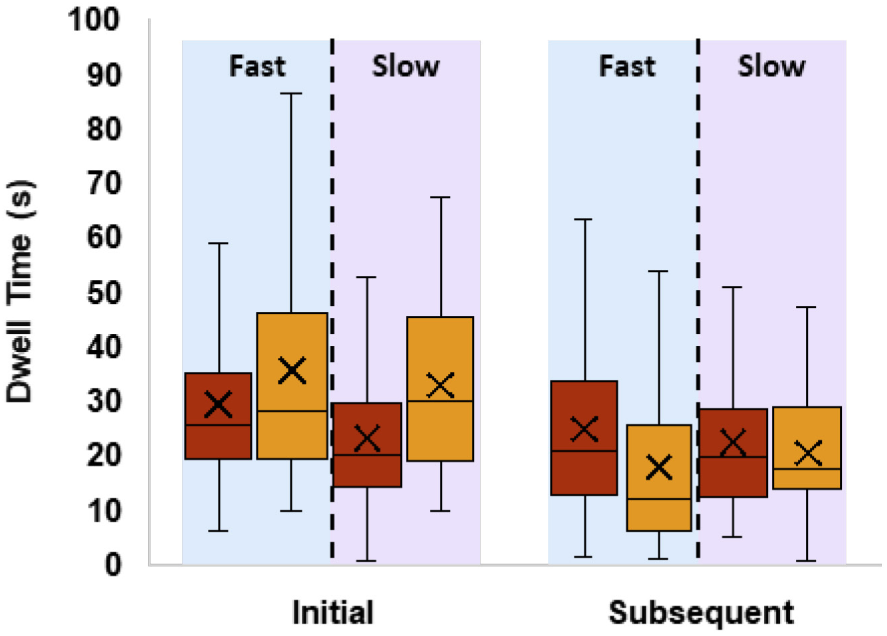
Initial and subsequent dwell times distribution based on construct and rate subpopulation for WT-MR (orange) and L101D-MR (red) of following transitions. The initial dwell time represents the time for a complex to productively assemble on a blunt end of dsDNA. The subsequent dwell times likely represent the time for productive assembly on recessed DNA ends. For each population, X indicates the average of the sample.

The L101D mutation within the MR complex decreases the initial dwell time before the first transition within a trace (post hoc, p<.0001, Fig. 5 and Supplemental Table 5). This is consistent with the pre-steady state kinetics indicating that the L101D-MR complex completes the productive-binding step more efficiently than the WT-MR complex^18^. This effect is observed for both slow (23.1 seconds compared to 33.4 second L101D- and WT-MR, respectively) and fast (29.7 seconds compared to 35.6 second L101D- and WT-MR, respectively) exonuclease events. Interestingly, although the slow WT-MR subpopulation also contains a disengaged dimer interface, this population does not display the shortened initial dwell times to the same extent as the L101D mutation, suggesting that L101D confers a more pronounced enhancement of productive binding.

The initial dwell time represents the time required for initial binding and productive assembly on blunt-ended dsDNA. Subsequent transitions and dwell times within a trace, when applicable, could be the result of release and rebinding of the recessed dsDNA end or a complex which pauses while bound to the DNA and resumes activity. These successive transitions accounted for 15 percent of WT-MR total transitions and 9 percent of L101D-MR total transitions. The observed difference between the proteins may reflect the more processive and slower exonuclease activity of L101D-MR, resulting in fewer intermediate substrates and thus fewer opportunities for subsequent exonuclease events. When comparing the initial and subsequent dwell times, subsequent dwell times are shorter in length than the initial dwell time across the different constructs and rate subpopulations (ANOVA, p<.0001). This is most apparent with the WT-MR complex leading up to both fast and slow transitions whereas the L101D complex shows a smaller difference between the initial and subsequent dwell times regardless of the type of transition that follows, indicating that the DNA end structure has less of an impact on L101D assembly efficiency (Fig. 5). Within the subsequent dwell times, the time leading up to WT-MR transitions is shorter than for L101D-MR (post hoc, p<.0001). This suggests that productive assembly for the WT-MR complex is more efficient on a recessed 3’ dsDNA substrate as compared to a blunt dsDNA end. The increased efficiency in productive assembly for WT-MR on recessed DNA substrates might indicate a positive feedback mechanism which promotes continued nucleolytic degradation which is not as prominent for L101D-MR. While subsequent dwell time data skewed toward the shorter duration dwell times, the WT-MR fast population had a severe skew that indicated a large number of dwell times that were likely caused by a complex pausing on the substrate briefly rather than complete DNA release followed by another MR complex binding (see Fig. 5 and Supplemental Fig. S7). It should be noted that subsequent dwell time was also independent of preceding transition rate and the length of the overhang (Supplemental Fig. S8 and S9). No determinable significant difference in length was observed between the second and third dwell times for either construct or rate subpopulation.

About two percent of transitions occurred immediately after the preceding transition (dwell time of zero). These transitions were the result of a sudden and notable change in rate at the end of a transition, moving a transition from one rate subpopulation to the other. In other words, a transition from a fast exonuclease event to a slow exonuclease event or vice versa. These transitions were treated separately from the other subsequent transitions because these observations are likely caused by major conformational changes within the complex and not rebinding events that make up the majority of the subsequent transitions. No notable trends in the rate and amplitude of these transitions were observed, although this data subset was likely too small to draw any statistically significant conclusions from. There was also no obvious trend favoring a slow transition converting to a fast transition more than a fast converting to a slow or vice versa.

### Impact of enzyme concentration on rate, amplitude, and dwell times

While studying the activity of an enzyme at a single concentration can provide valuable information on the kinetics of the complex, there are some limitations that can best be overcome by observing the single molecule assays at varying concentrations. In particular, we were interested in the concentration dependence of rate, amplitude, and dwell times which could indicate how many enzyme complexes contributed to a single observed exonuclease event and whether subsequent events in a single trace were the result of new binding events or if the same complex was paused on the DNA substrate.

The rate of nucleotide excision appears to be independent of concentration within the constraints of this study (Supplemental Fig. S10). We did, however, observe a slight positive correlation between the amplitude of the transitions and enzyme concentration for L101D-MR (Fig. 6). This trend suggests that multiple L101D-MR complexes might contribute to a single transition. One possible interpretation is that a secondary complex waits to resume activity once the initiating enzyme complex disengages. However, our data does not show an increase in transitions featuring sudden rate changes — previously described as a dwell time of zero between two transitions with different rates (Supplemental Table S6) — alongside the increasing processivity. If we are not seeing an increase in intra-transition rate changes in correlation to an increase in complexes contributing to each transition, then the initiating complex rate must be determinate of the entire transition’s rate. Rather than the initial complex releasing and making way for the secondary complex to take over, this indicates that the initial enzyme complex determines the exonuclease rate, while the secondary complex may promote higher processivity by binding to the ssDNA product directly behind the initiating complex and reducing the potential for backtracking by the initiating enzyme complex.

**Figure 6.**
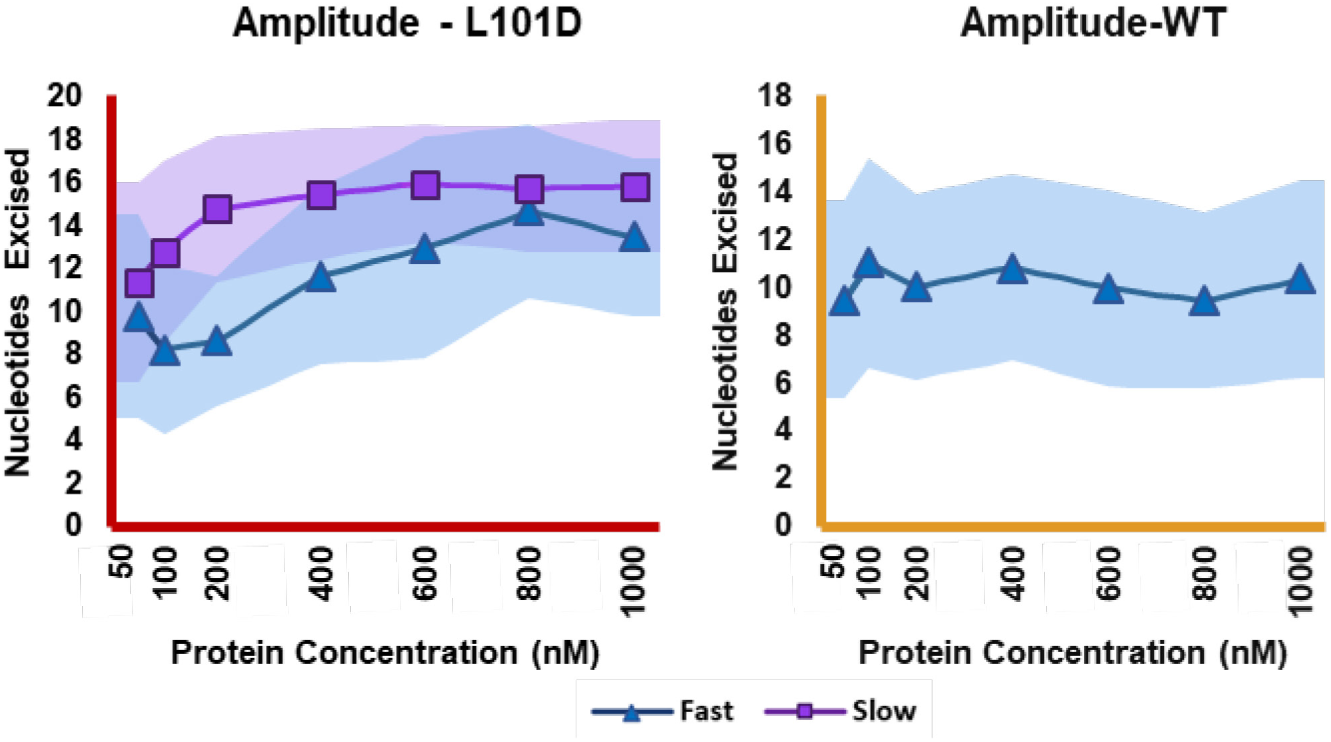
Average number of nucleotides excised per exonuclease event by concentration. Fast (blue) and slow (purple) rate subpopulations are shown separately with the shaded regions representing one standard deviation of the subpopulation. There was an insufficient number of WT slow transitions to include in analysis.

Although this subtle trend was consistently observed for both fast and slow exonuclease events in the L101D-MR complexes, it was absent in both fast and slow WT-MR populations - although the sample size for the slow-rate WT-MR subpopulation is too small to draw statistically significant conclusions. The dissociation constants for ssDNA and dsDNA binding are nearly equivalent between WT-MR and L101D-MR, making it unlikely that binding affinity explains this phenomenon^18^. One contributing factor could be the rate at which WT-MR completes its activity. Even when compared to the L101D-MR fast subpopulation, the WT-MR fast subpopulation still has an average rate at least 2x greater (Figure 3c). Because of the previously discussed maximum rate limitation, this particular data subset likely underestimates the WT-MR fast rate, making its difference from the L101D-MR fast rate potentially even greater. This high rate could limit the opportunity for another complex to bind and noticeably impact the amplitude of these transitions. Another reasonable hypothesis would be that the engaged dimer interface of the WT-MR construct reduces the rate of backtracking that is observed in other conformations such as the disengaged dimer interface of the slow populations and the, perhaps slightly altered, dimer interface of fast L101D-MR subpopulation.

As expected, initial dwell times decreased with increasing enzyme concentration (Fig. 7). This trend was most noticeable with WT-MR over the range tested (50 nM to 1000 nM) with saturation being reached by 600 nM. L101D-MR, which in all instances tended to initiate nuclease events faster that WT-MR, seemed to reach saturation by 200 nM enzyme concentration. This variation is also likely caused by the competitive, nonproductive binding that is suspected to be found within the WT population.

**Figure 7.**
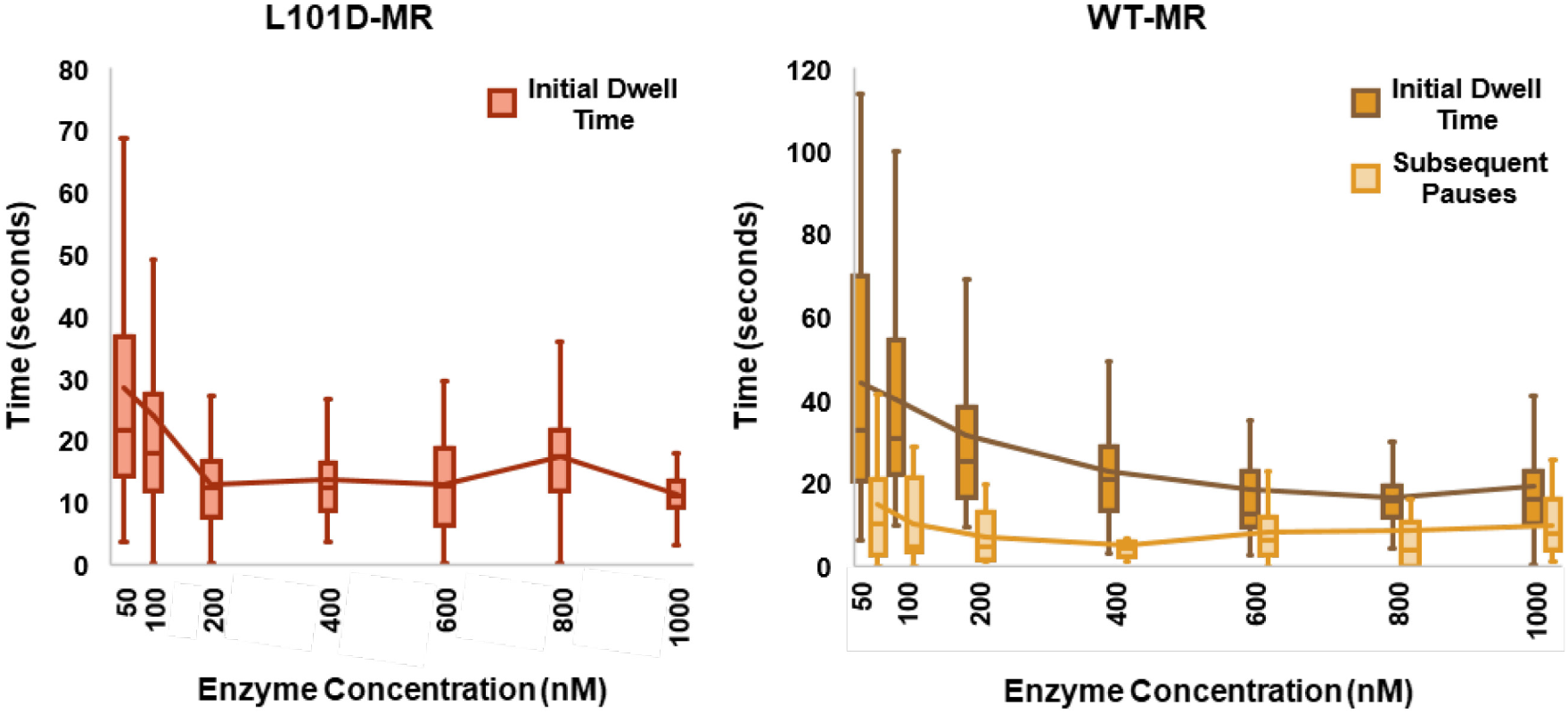
Initial and subsequent dwell times for L101D-MR (red) and WT-MR (orange). For WT-MR, subsequent dwell times (light orange) remain mostly unchanged across the different concentrations. Initial dwell times for WT-MR (dark orange) show a decrease in duration across the different concentrations. A slight decrease is observed in the lowest concentrations for L101D-MR for the initial dwell time (red) as well. There was insufficient data of the subsequent dwell times for WT-MR to include in analysis. The line follows the average across the different concentrations.

While our data set for subsequent dwell times is more limited across the various concentrations, we still observed that WT-MR generally displays shorter subsequent dwell times than initial dwell times. Over the concentration range tested, these subsequent dwell times appear independent of concentration (averaging ∼11.4 seconds for WT-MR), suggesting that enzyme concentration becomes rate-limiting only below 50 nM under our conditions. Additionally, a substantial fraction of these subsequent WT-MR dwell times likely reflects fast-subpopulation complexes pausing briefly on the DNA substrate rather than new binding events. This interpretation is supported by Figure 5 (and Supplemental Figure S7), where the WT-MR-fast subpopulation shows a pronounced skew toward shorter subsequent dwell times compared to other subpopulations.

**Figure 8.**
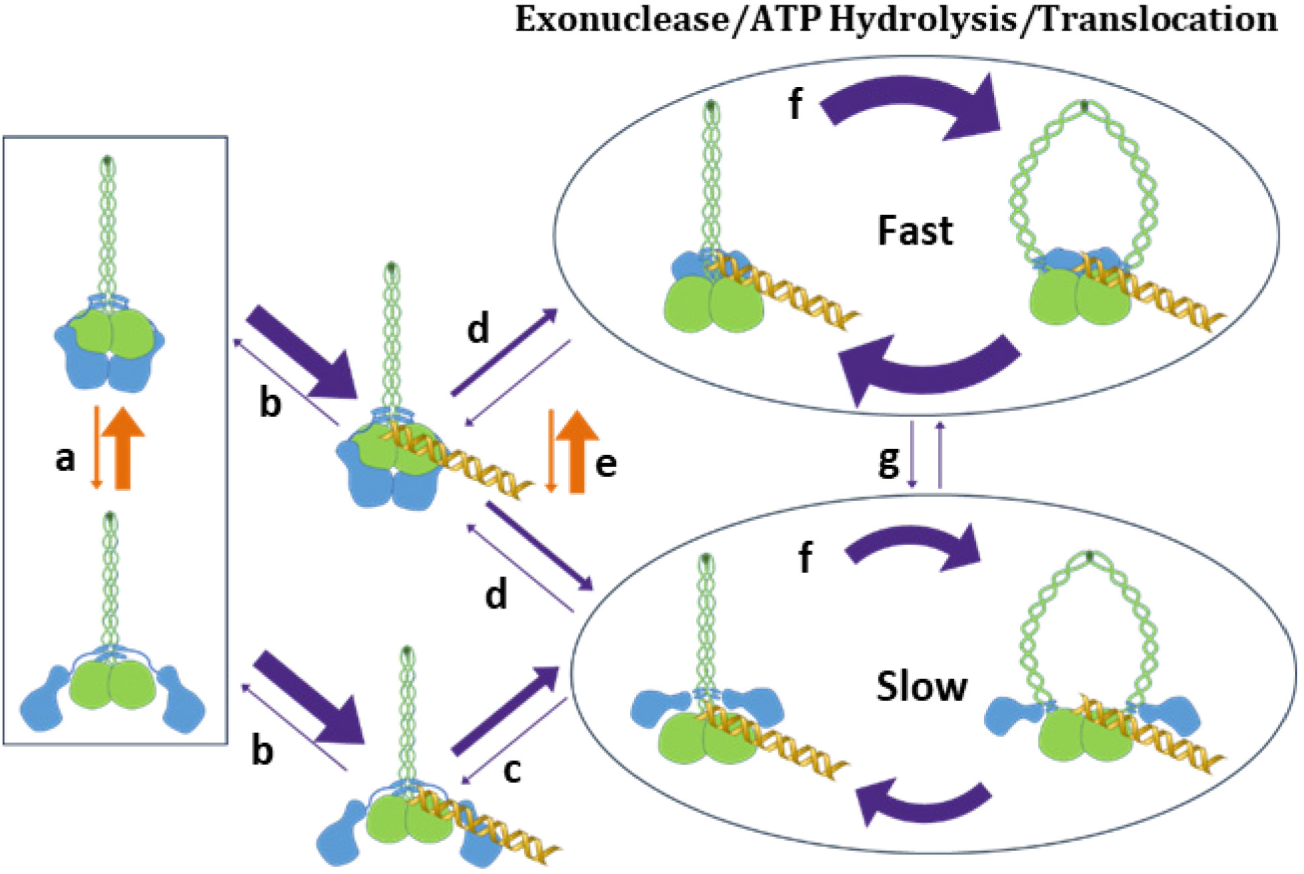
Mechanism of nuclease activity for T4-Phage, WT-MR on blunt-end dsDNA. Orange arrows (a and e) indicate equilibriums. Dark purple arrows indicate rates. a) Negative Stain EM images indicate the complex exists in both the disengaged and engaged configurations in the absence of DNA. Due to the higher volume of fast exonuclease events, it seems that the equilibrium likely lies in the direction favoring the engaged dimer interface for WT-MR. The equilibrium lies heavily in the direction of the open dimer interface for L101D-MR. b) Previous ensemble assays comparing L101D-MR and WT-MR indicate that constructs with the disengaged and engaged dimer interface have the same binding affinity for dsDNA. c) Data from L101D indicate that constructs with disengaged dimer interfaces will productively assemble at a faster rate than those with engaged dimer interfaces. d) Due to similar rates of productive assembly between WT fast and slow populations, it appears that the majority of slow WT exonuclease events result from the dimer interface falling apart upon initiation of nuclease activity. Productive assembly is the rate-limiting step for both L101D- and WT-MR. e) The majority of constructs remain in the engaged conformation upon initiation of exonuclease activity. f) During activity, the shift between the nuclease state and translocation state occurs at a much faster rate for constructs with an engaged dimer interface. g) During activity, a conformational change from disengaged to engaged or vice versa occurs less than 2 % of the time.

## DISCUSSION

Previous mechanistic models of the MR complex in the presence of ATP and dsDNA have suggested a rate-limiting productive assembly step followed by fast and processive exonuclease activity^16,18^. While ATP is not necessary for the excision of the first nucleotide, ATP hydrolysis is necessary for driving translocation across the ssDNA product and, in turn, the continuation of exonucleolytic substrate degradation^16^. Our study of the MR complex using smFRET has revealed a 2^nd^ mechanism of nuclease activity which is characterized by a slower, yet more processive, translocation that is mediated by a form of the MR complex with a disengaged Mre11 dimer interface. Further investigation of this subpopulation has provided a deeper insight into the structural basis of MR complex function.

Previous studies have proposed that exonuclease activity involves cycling of the complex between nuclease and translocation conformational states^18^. Our results indicate that, independent of exonuclease rate, the WT-MR complex transitions more efficiently between these two states compared to the L101D-MR complex. Previous studies have indicated that L101D-MR’s intrinsic nuclease activity is equivalent to WT-MR and that the L101D-MR complex shows minimal activation of ATPase activity in the presence of DNA, indicating that the disengaged dimer interface primarily affects the translocation state^18^. It was originally suggested that ATP hydrolysis causes the L101D-MR complex to release DNA. Our data, however, indicates that ATP hydrolysis is still necessary to drive translocation of the L101D-MR complex and that the reduction in activity is due to a reduced rate in translocation and not reduced processivity.

The results indicate that WT-MR has lower processivity than L101D-MR, at least on the relatively short DNA substrate used in this study. This may indicate that the disengaged dimer interface of L101D-MR creates less hindrance for DNA substrates attempting to reach the nuclease active site of Mre11 or a regulatory domain at the dimer interface that hinders processivity is absent. The WT-MR complex has an increased number of nonproductive binding events, which could indicate that Mre11, when lacking the physical restrictions of an engaged dimer interface, has more flexibility to maneuver around a DNA end that is not optimally positioned with respect to the complex which, in turn, reduces nonproductive binding events.

The differences between the conformations of the L101D-MR and WT-MR complexes for both the engaged and disengaged MR subpopulations, which contribute to this variation in activity, likely also contributes to the variations we see at the rate-limiting productive assembly step. During the initiation of exonuclease activity, we observe a faster rate of productive assembly for L101D-MR compared to WT-MR, with the WT-MR complex showing minimal variation in assembly rates across the different rate subpopulations. This suggests that the WT-MR complex disfavors the disengaged dimer interface when not bound to DNA and that the slow-rate subpopulation we observe is due to interface disruption upon initiation of exonuclease activity. Perhaps there is an in vivo advantage to the Mre11 dimer interface separating readily if the conformation of the DNA bound to the complex is not optimal for DNA processing. This indicates a trade-off in which the MR complex sacrifices speed in favor of improved interaction with complex DNA conformations, including secondary structures or blocked ends. The possibility of the Mre11 dimer interface disengaging upon DNA binding and not while performing exonuclease activity is consistent with previous kinetic studies indicating the rate limiting productive assembly step is likely caused by a major conformational change that only occurs during initiation and not throughout the reaction^24^. Cryo-EM structures indicate that Mre11 moves from a position distal to the coiled-coil domain up next to the coiled-coil domain upon DNA binding^31^.

The apparent advantage of the disengaged dimer interface in hastening productive assembly on a blunt DNA end is diminished when binding to a recessed end as indicated by approximately equivalent dwell times for both L101D and WT leading up to 2^nd^ and 3^rd^ transitions. While the initial dwell time for WT was greater than the subsequent dwell times for both rate subpopulations, for L101D-MR the duration of the initial dwell times of the slow rate subpopulation and the subsequent dwell times for both fast and slow subpopulations were nearly identical. This would support the idea that a large portion of the L101D-MR population is in the disengaged conformation prior to interacting with DNA and that this disengaged dimer interface upon initial binding allows for bypassing a regulatory mechanism that imposes stricter requirements on the DNA substrate.

We observe that both L101D-MR and WT-MR populations can be divided into two subpopulations based on exonuclease activity rates, with the majority of WT-MR transitions exhibiting fast activity and the majority of L101D-MR transitions exhibiting slow activity. Interestingly, we observed an apparent higher processivity for transitions of slow exonuclease activity compared to those with faster activity. Our data also demonstrates that L101D-MR is processive and is primarily affected by reduced translocation. However, the positive correlation between nucleotides excised and enzyme concentration indicates that more than one enzyme complex may contribute to a single L101D-MR transition. We have found that the smFRET kinetics analysis is consistent with ensemble assays: when the Mre11 dimer interface is disrupted by the L101D mutation, pause times decrease compared to WT-MR, suggesting that the rate-limiting assembly step becomes more efficient.

Building on our findings of dual exonuclease subpopulations and the impact of the Mre11 dimer interface, future investigations will focus on characterizing MR complex conformational changes at the single-molecule level. Such studies will clarify how different conformations emerge during ATP hydrolysis, the initiation of nuclease activity, and the excision of individual nucleotides, leading to a more comprehensive mechanistic understanding of MR-mediated DNA repair.

## ASSOCIATED CONTENT

### Supporting Information

The Supporting Information is available free of charge at ___:

- Analytical methods, tables with statistical analyses, tables with counts of relevant populations, additional figures obtained from FRET data, supporting data from ensemble assays and negative-stain EM images

## AUTHOR INFORMATION

### Author Contributions

S.W.N., J.B.C., and T.T. conceived of and planned the study. Y.G. performed the negative stain EM. J.B.C. and T.T. performed all other experiments. J.B.C. processed and analyzed the data.

S.W.N. acquired funding and supervised the project. T.T. and J.B.C. wrote the original draft, and J.B.C. and S.W.N. edited the draft.

### Notes

The authors declare no competing financial interest.

## ACKNOWLEDGEMENTS

This work was financially supported by National Science Foundation Grant MCB:1121693.

## Notes

### Competing Interest Statement

The authors have declared no competing interest.

