## Supplemental Figures for "Single-Molecule FRET Reveals Two T4 Phage MR Complex Exonuclease States Regulated by the Mre11 Dimer Interface"

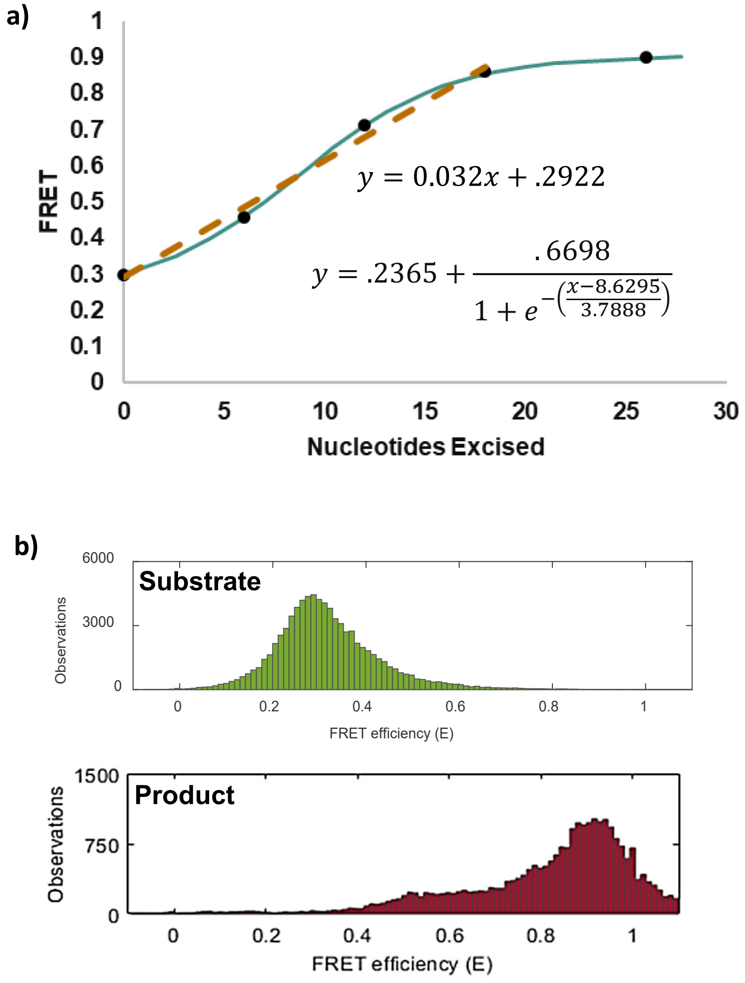


**Figure S1: Conversion from FRET to nucleotides excised.** a) A slight sigmoidal shape was observed with the curve plateauing between 18 nucleotides excised and 26 nucleotides excised. Shown in teal, the fitted sigmoidal curve and, shown in brown, the linear conversion that was used in the calculations of this study. The linear and sigmoidal equations used are shown. The black dots are the raw data points. b) The raw data points came from fitting a gaussian curve to the distribution of FRET values across two-minute movies of the given product mimic and taking the FRET value at the apex of the curve. In green, the original substrate (no nucleotides excised) and, in red, the complete product (26 nucleotides excised). The example histograms show the count of frames from all traces from one video which fall into designated FRET bins. Two to three movies from different days were used for each product mimic.

**Figure S2: Efficiency of exonuclease activity on Cy3/Cy5-labeled DNA and 2-AP-labeled DNA**

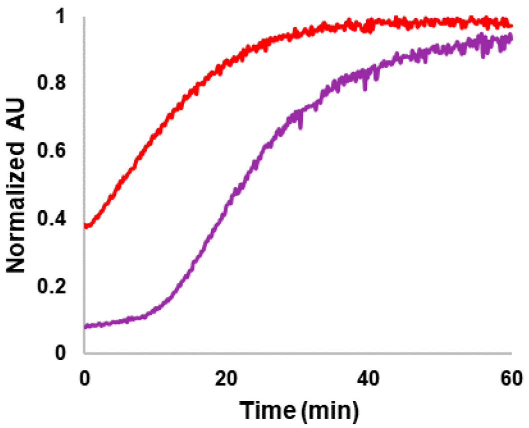


To determine if the DNA substrate we intended to use for the single-molecule experiments replicated our observed rates in bulk assays, we directly compared the Cy3/Cy5 labeled substrate with our standard 2-aminopurine (2-AP) DNA substrate. The standard DNA substrate is 50 base-pairs long with the 2-AP label located at the $22^{nd}$ nucleotide from the 3' terminal. On the fluorescent substrate, Cy3 and Cy5 were positioned 20 base pairs apart. In the reactions (data shown to the right), 1 uM of DNA substrate was used with 0.2 uM of the WT-MR complex.

The data indicates that the Cy3 and Cy5 dyes do not interfere with the exonuclease activity of the MR complex. Moreover, the Cy3/Cy5 substrate displayed a possible advantage over 2-AP assays. The fluorescence intensity from the 2-AP DNA substrate with the label at the $22^{nd}$ nucleotide position shows only a minor increase until enough degradation has taken place to release most of the fluorescent labels from the substrate (i.e., it is an all-or-nothing assay where the signal only increases if 22 nucleotides are excised). In contrast, Cy3/Cy5–labeled DNA is sensitive to changes caused by DNA degradation starting from earliest time points, allowing for observation of all reaction intermediates.

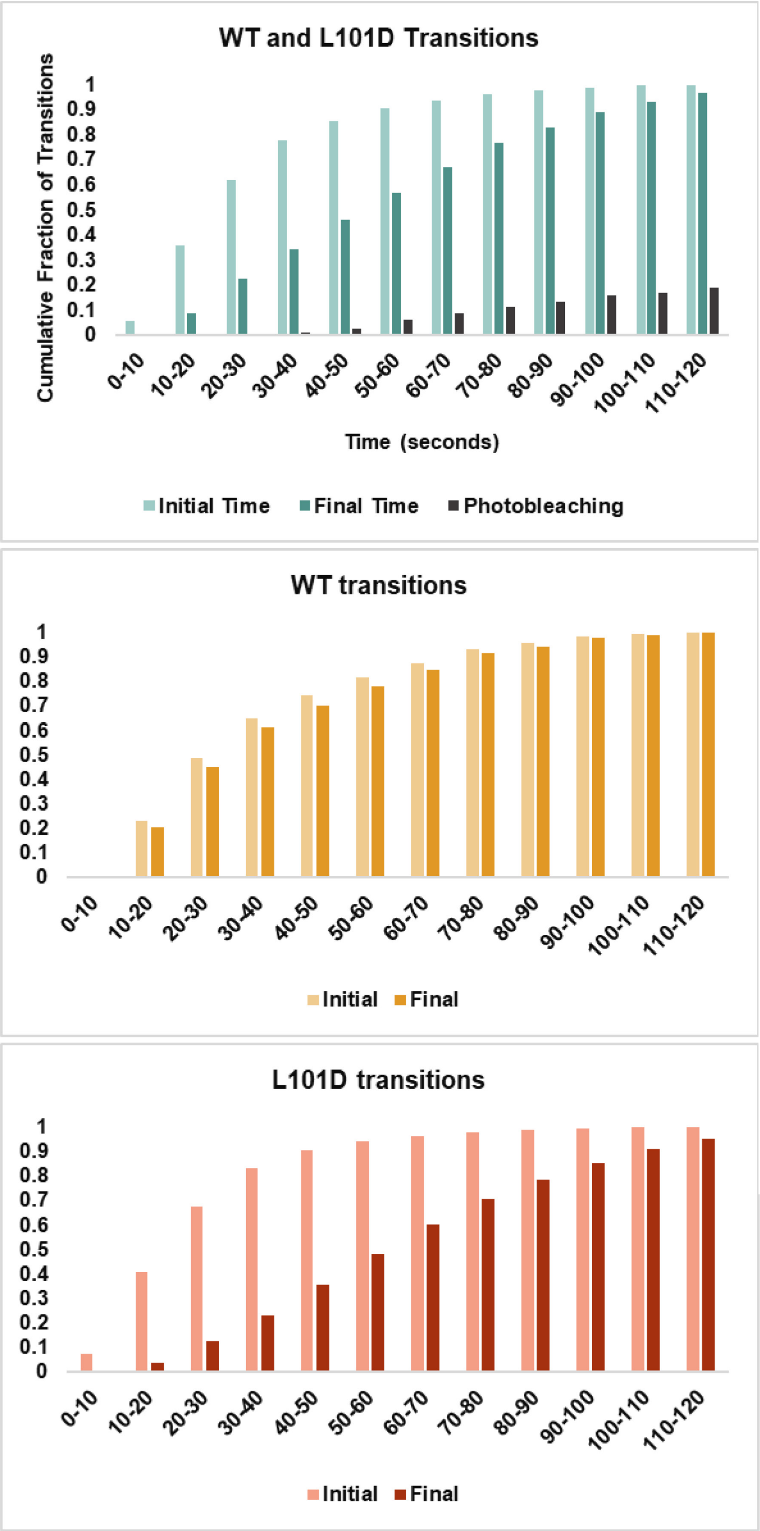


**Figure S3: Photobleaching over time in comparison with the initiation and completion of transitions.** Here, the estimated amount of photobleaching over time is compared to the number of traces initiating and finishing over 2 minutes. The accumulation over time of all transitions (top, teal) initiated (light teal) and completed (dark teal) compared to the increase with time of photobleaching. Less than 20% of all traces (including complete transitions, incomplete transitions, and traces with uninitiated activity) photobleach by the end of two minutes. WT-MR (middle, red) and L101D-MR (bottom, orange) divided out to show the accumulation of initiated (light) vs completed (dark) transitions over time. For both WT-MR and L101D-MR, the number of transitions initiated has plateaued and the number of transitions reaching completion is nearly equivalent in number by the end of the two-minute movie.

a)

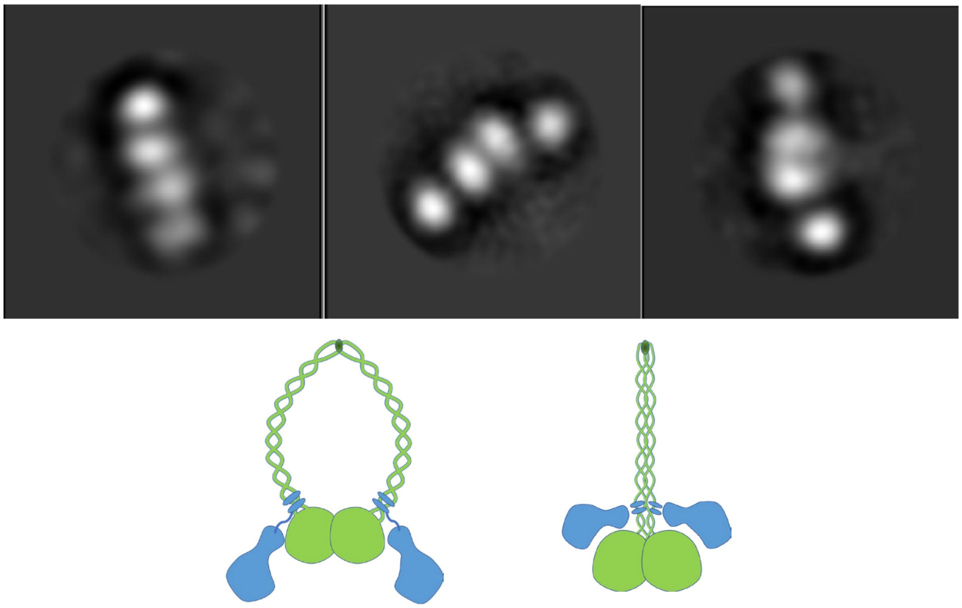

b)

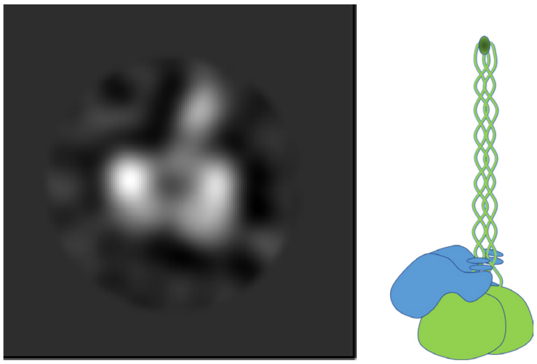

**Figure S4: Negative-stain transmission EM images of WT-MR in the presence of ATP$\gamma$S and the absence of DNA.** a) Representative 2D classifications of WT-MR complex in the extended conformation. Based on earlier structure information of MR complex, we suspect the center dimer represent Rad50 dimer whereas the separated density at the sides are Mre11 subunits. The coiled-coil domains are difficult to distinguish with confidence in these 2D averages. b) An example of 2D class image capturing the MR complex possibly in a compact conformation. The dimer density appears larger than that in panel an and we thus suspect that the density may represent MR dimer. In addition, part of the coiled-coil domain may be visible in this conformation. The images were collected on a Technai 12 electron microscope (FEI) operated at 120 kV and 300,000 magnification. The CTF estimation and 2D classification were performed with RELION 3.0[1].

(1) Zivanov, J.; Nakane, T.; Forsberg, B. O.; Kimanius, D.; Hagen, W. J.; Lindahl, E.; Scheres, S. H. New Tools for Automated High-Resolution Cryo-EM Structure Determination in RELION-3. *Elife* **2018**, *7*, e42166. https://doi.org/10.7554/eLife.42166.

**Figure S5: Rad50-independent exonuclease activity**

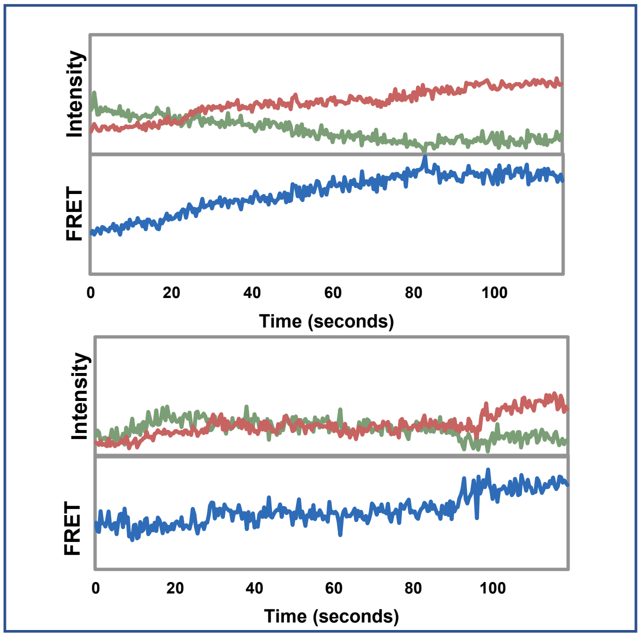


The deletion of the Rad50 binding domain of T4-Mre11, along with the flexible linker between the globular and the C-terminal domains (ΔC289), increases the mutant protein's DNA binding affinity. This leads to an increase in exonuclease activity that can be detected at much lower concentrations compared to the WT-Mre11. Here, 4uM of ΔC289 has been used as a surrogate for WT-Mre11 and likely represents the type of exonuclease activity that would be observed in WT-Mre11 reactions. For the two example sets of traces, the fluorescent intensity of Cy3 (green) and Cy5 (red) is shown over time along with the calculated FRET trace (blue). The data indicate that the majority of the FRET transitions for ΔC289 show a processive, gradual increase FRET. A small number of single-step rises are observed, but these are of very low amplitude.

**Figure S6: Investigation of possible hidden events**

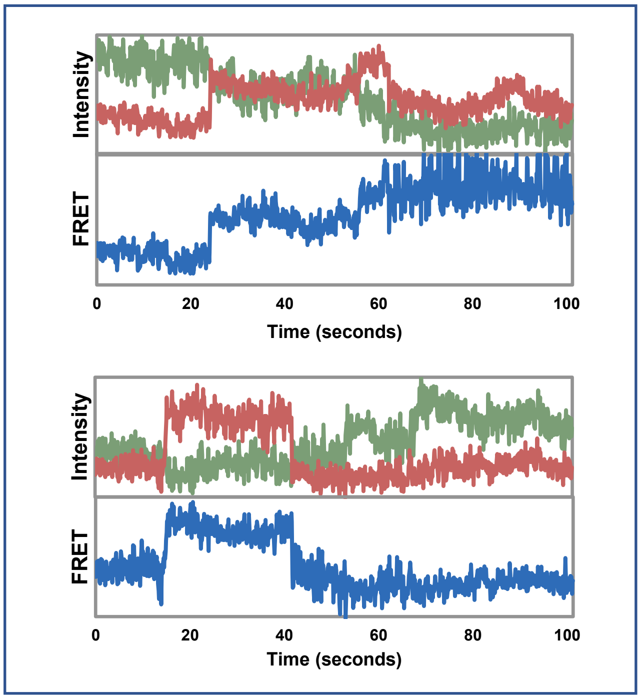


A diverse set of changes in biomolecular processes occurring on a time scale of ~10 ms or slower have been studied using sm-FRET. However, achieving a high signal-to-noise ratio when imaging at faster exposure (sampling) times requires high intensity laser excitation, which favors photobleaching and shortens the usable observation time. While 400 ms exposure time has been used for most of the fluorescence imaging, preliminary data from imaging done at 100 ms exposure time suggests that the gradual and single-step FRET transition patterns remain essentially unchanged (right). For the three example sets of traces, the fluorescent intensity of Cy3 (green) and Cy5 (red) is shown over time along with the calculated FRET trace (blue).

Fast Transitions

| | SS | Df | F value | p value |
|---|---|---|---|---|
| (Intercept) | 68039 | 1 | 582 | < 2.2e-16 *** |
| Construct | 22086 | 1 | 188.9 | < 2.2e-16 *** |
| Residuals | 2E+05 | 1349 | | |

Slow Transitions

| | SS | Df | F value | p value |
|---|---|---|---|---|
| (Intercept) | 230 | 1 | 3619 | < 2.2e-16 *** |
| Construct | 1.013 | 1 | 15.95 | 6.774e-05 *** |
| Residuals | 114.6 | 1804 | | |

**Table S1: 1-way ANOVA type III tests** were run on linearized models of the fast and slow subpopulations separately to assess the variance of rate between the constructs (WT-MR and L101D-MR). Post hoc analysis was deemed unnecessary due to the nature of the ANOVA and the data.

**Figure 3c: Rate population cutoff determination**

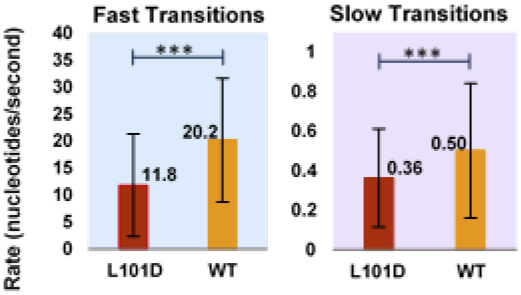


**Transition Counts**

| | fast | slow |
|---|---|---|
| **L101D** | 490 | 1752 |
| **WT** | 861 | 54 |

| Variables | SS | Df | F value | p value |
|---|---|---|---|---|
| (Intercept) | 25715 | 1 | 1375.153 | <2.20E-16 |
| construct | 2025 | 1 | 108.294 | <2.20E-16 |
| rate | 10731 | 1 | 573.841 | <2.20E-16 |
| construct:rate | 939 | 1 | 50.222 | 1.69E-12 |
| Residuals | 58961 | 3153 | | |

Response: nucleotides excised

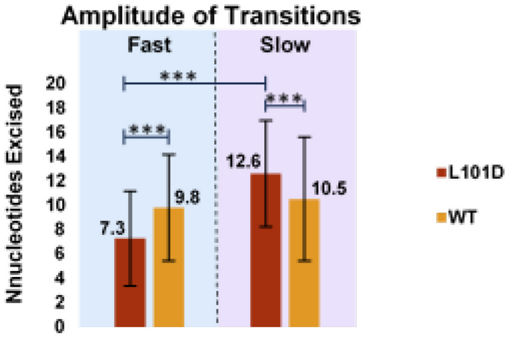


**Table S2: ANOVA Type III test for amplitude analysis (Figure 4)**
Significance was determined using R and R Studio. Because the data was unbalanced, a linearized model was run in correlation to a 2-way ANOVA with type III sums of squares. Both of the main effects (construct and rate) had a significant impact on the number of nucleotides excised as well as the interaction between construct and rate.

**Dependent Variable**

| construct | rate | mean | SE | df | lower.CL | upper.CL |
|---|---|---|---|---|---|---|
| L101D | fast | 7.24 | 0.195 | 3153 | 6.86 | 7.63 |
| WT | fast | 9.79 | 0.147 | 3153 | 9.5 | 10.08 |
| L101D | slow | 12.54 | 0.103 | 3153 | 12.34 | 12.74 |
| WT | slow | 10.51 | 0.588 | 3153 | 9.36 | 11.66 |
| | fast | 8.52 | 0.122 | 3153 | 8.28 | 8.76 |
| | slow | 11.52 | 0.299 | 3153 | 10.94 | 12.11 |
| L101D | | 9.89 | 0.11 | 3153 | 9.67 | 10.1 |
| WT | | 10.15 | 0.303 | 3153 | 9.56 | 10.7 |

**Transition Counts**

| | fast | slow |
|---|---|---|
| **L101D** | 490 | 1752 |
| **WT** | 861 | 54 |

| variable/s | constant | contrast | estimate | SE | df | t.ratio | p value |
|---|---|---|---|---|---|---|---|
| C:R | fast | L101D-WT | -2.55 | 0.245 | 3153 | -10.406 | <.0001 |
| C:R | slow | L101D-WT | 2.03 | 0.597 | 3153 | 3.396 | 0.0007 |
| C:R | L101D | fast-slow | -5.294 | 0.221 | 3153 | -23.955 | <.0001 |
| C:R | WT | fast-slow | -0.718 | 0.607 | 3153 | -1.184 | 0.2365 |
| R* | | fast-slow | -3.01 | 0.323 | 3153 | -9.312 | <.0001 |
| C* | | L101D-WT | -0.259 | 0.323 | 3153 | -0.802 | 0.4228 |

*Results may be misleading due to involvement in interactions

**Table S3: Tukey's HSD Post Hoc Test following ANOVA (Table Sw)**

Following ANOVA, the Post hoc test indicated that there was not a significant difference between the fast and slow subpopulations for WT-MR (p>.05).

### Counts of Dwell Times

| | WT | | L101D | |
|---|---|---|---|---|
| | fast | slow | fast | slow |
| Initial | 724 | 37 | 368 | 1627 |
| Sub. | 133 | 6 | 115 | 77 |

| Variables | SS | Df | F value | p value | |
|---|---|---|---|---|---|
| (Intercept) | 127.398 | 1 | 1771.59 | < 2.2e-16 | *** |
| construct | 3.752 | 1 | 52.1765 | 6.37E-13 | *** |
| step | 1.555 | 1 | 21.6169 | 3.47E-06 | *** |
| rate | 1.498 | 1 | 20.83 | 5.22E-06 | *** |
| construct:step | 4.077 | 1 | 56.7 | 6.63E-14 | *** |
| step:rate | 0.437 | 1 | 6.0795 | 0.01373 | * |
| construct:rate | 0.336 | 1 | 4.6670 | 0.03082 | * |
| Residuals | 221.488 | 3080 | | | |

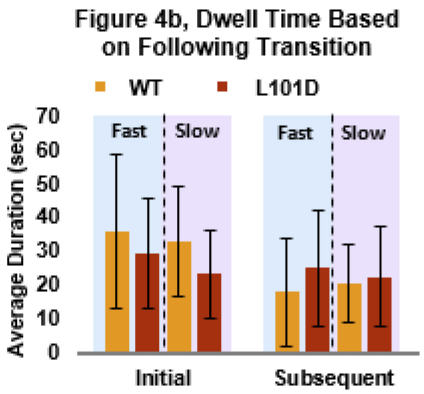


**Table S4: ANOVA (Type III Tests) for dwell time analysis (Figure 5)**
Significance was determined using R and R Studio. Because the data was unbalanced, a linearized model was run in correlation to a 3-way ANOVA with type III sums of squares. Prior to running ANOVA, the optimal model was determined comparing BIC, AIC, and F-tests of all possible models. The optimal model included all three main effects and three interacting terms (construct:step, step:rate, and rate:construct). A Tukey HSD test was performed following ANOVA (below).

**Dependent Variables**

| rate | step | construct | mean | SE | df | lower.CL | upper.CL |
|---|---|---|---|---|---|---|---|
| | Initial | L101D | 1.35 | 0.00772 | 3080 | 1.34 | 1.37 |
| | Initial | WT | 1.46 | 0.02142 | 3080 | 1.42 | 1.50 |
| | Subsequent | L101D | 1.27 | 0.01971 | 3080 | 1.23 | 1.31 |
| | Subsequent | WT | 1.11 | 0.03363 | 3080 | 1.04 | 1.17 |
| | Initial | | 1.41 | 0.0113 | 3080 | 1.39 | 1.43 |
| | Subsequent | | 1.19 | 0.0202 | 3080 | 1.15 | 1.23 |
| Fast | | | 1.31 | 0.00955 | 3080 | 1.29 | 1.33 |
| Slow | | | 1.29 | 0.02474 | 3080 | 1.24 | 1.34 |
| | | L101D | 1.31 | 0.0106 | 3080 | 1.29 | 1.33 |
| | | WT | 1.29 | 0.0236 | 3080 | 1.24 | 1.33 |
| Fast | Initial | | 1.44 | 0.00858 | 3080 | 1.43 | 1.46 |
| Slow | Initial | | 1.37 | 0.02102 | 3080 | 1.33 | 1.41 |
| Fast | Subsequent | | 1.18 | 0.01707 | 3080 | 1.14 | 1.21 |
| Slow | Subsequent | | 1.20 | 0.03694 | 3080 | 1.13 | 1.28 |
| Fast | | L101D | 1.35 | 0.0142 | 3080 | 1.32 | 1.38 |
| Fast | | WT | 1.27 | 0.0125 | 3080 | 1.25 | 1.30 |
| Slow | | L101D | 1.28 | 0.0152 | 3080 | 1.25 | 1.31 |
| Slow | | WT | 1.30 | 0.0441 | 3080 | 1.21 | 1.38 |

| variable/s | constant | contrast | estimate | SE | df | t.ratio | p value |
|---|---|---|---|---|---|---|---|
| S | | Init.-Sub. | 0.218 | 0.0191 | 3080 | 11.408 | <.0001 |
| R | | fast - slow | 0.0213 | 0.0265 | 3080 | 0.803 | 0.4219 |
| C | | L101D - WT | 0.0287 | 0.0252 | 3080 | 1.136 | 0.2562 |
| R:C | Fast | L101D - WT | 0.0765 | 0.0188 | 3080 | 4.07 | <.0001 |
| R:C | Slow | L101D - WT | -0.0192 | 0.0436 | 3080 | -0.441 | 0.6594 |
| C:R | L101D | fast - slow | 0.0692 | 0.0206 | 3080 | 3.362 | 0.0008 |
| C:R | WT | fast - slow | -0.0266 | 0.0443 | 3080 | -0.6 | 0.5486 |
| S:R | Initial | fast - slow | 0.0706 | 0.0228 | 3080 | 3.094 | 0.002 |
| S:R | Subsequent | fast - slow | -0.028 | 0.041 | 3080 | -0.683 | 0.4945 |
| R:S | Fast | Init.-Sub. | 0.267 | 0.0191 | 3080 | 13.989 | <.0001 |
| R:S | Slow | Init.-Sub. | 0.168 | 0.0341 | 3080 | 4.936 | <.0001 |
| C:S | L101D | Init.-Sub. | 0.0805 | 0.0212 | 3080 | 3.797 | 0.0001 |
| C:S | WT | Init.-Sub. | 0.3551 | 0.0307 | 3080 | 11.553 | <.0001 |
| S:C | Initial | L101D - WT | -0.109 | 0.023 | 3080 | -4.735 | <.0001 |
| S:C | Subsequent | L101D - WT | 0.166 | 0.0376 | 3080 | 4.418 | <.0001 |

**Table S5: Tukey HSD Post hoc test for dwell time analysis**

Rate and construct are only significant when all interactions are taken into consideration.

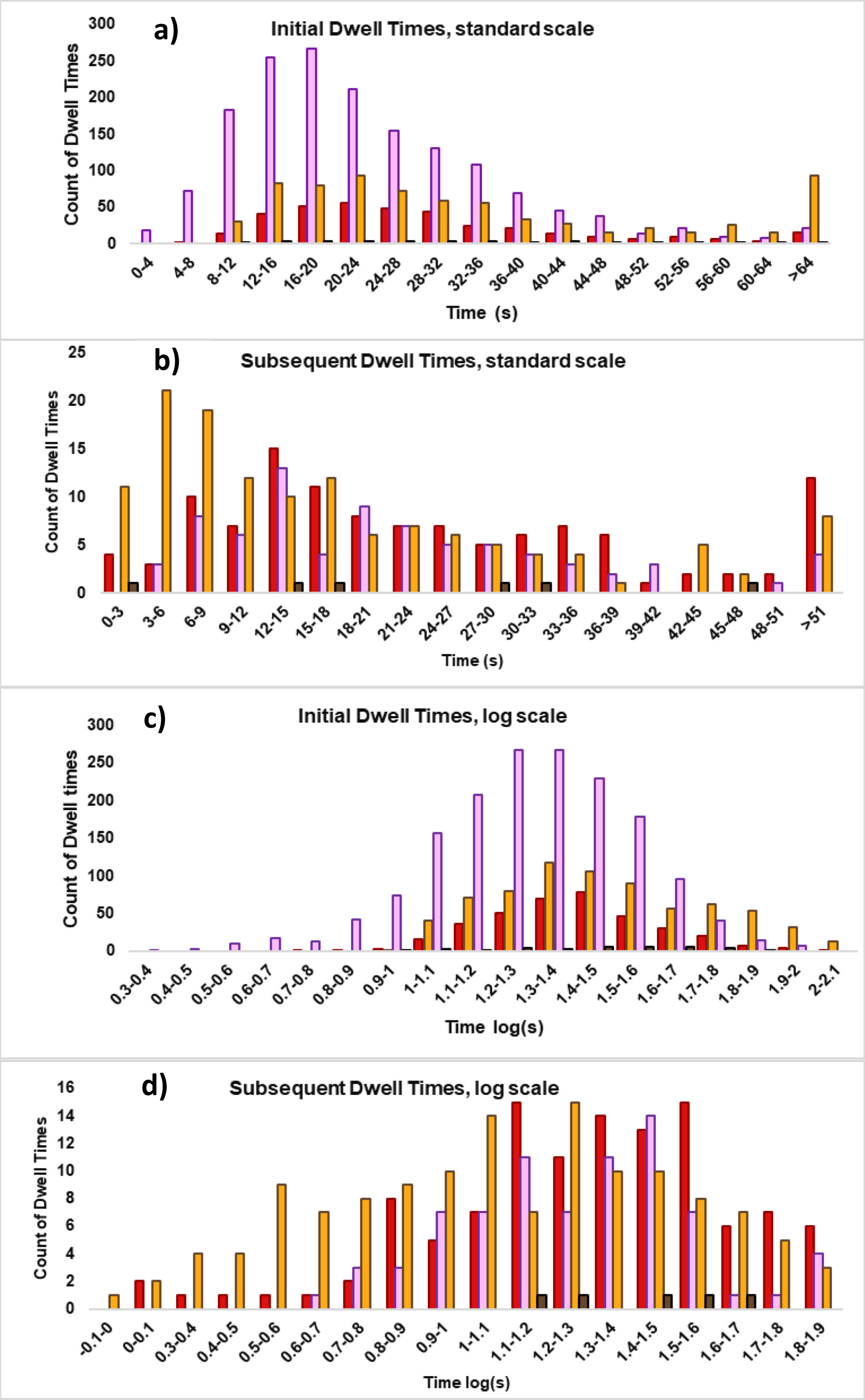


**Figure S7: Dwell time data from Figure 5** in both a standard scale and converted to a log scale for statistical analysis. WT-MR-fast (orange), L101D-MR-fast (red), WT-MR-slow (brown), L101D-MR-fast (pink) ANOVA, while fairly robust against skewed data (a and c), would be preferably run on normally distributed data. We achieved a more normal distribution by applying a logarithmic scale to the data set (b and d). The resulting analysis of the logarithmized data is what we are presenting here. However, a parallel analysis (not shown) of the raw data produced similar results in regards to statistically significant components of the model.

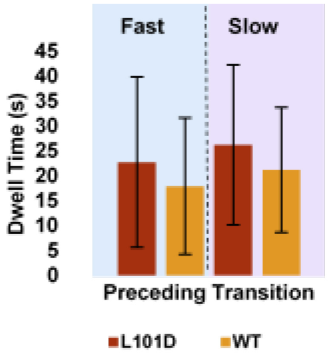


**Figure S8: Subsequent dwell time length in regards to the preceding transition rate.** Dwell times of zero (changes in rate) were not included in this data set. Preceding transition rate had no significant impact on dwell time (See statistical analysis Table S!).

| | SS | Df | F value | Pr(>F) |
|---|---|---|---|---|
| (Intercept) | 108485 | 1 | 420.51 | < 2.2e-16 *** |
| construct | 2680 | 1 | 10.39 | 0.001395 ** |
| Residuals | 84619 | 328 | | |

**Table S6: ANOVA analysis of dwell times in regards to preceding transition rate (Figure S8)**
For the data displayed in Figure S7 the main effects and interactions of construct and preceding transition rate subpopulation were modeled against a response variable of dwell time. After choosing the optimal linearized model by running BIC, AIC, and an F-test, the model of best fit was determined to only include the main effect of construct. This signifies that preceding rate had no siginficant impact on the duration of the dwell times. Because the construct will always be identical for the preceding and following transitions, it is not possible to know for certain the impact preceding construct has on dwell time. However, the dependency of dwell time on the construct within interactions with step and rate of following transitions (Figure 4b and statistical analysis in Table Sj and SS) confirms that at least a portion of the significance of construct in regards to dwell time lies solely with the following transition if not in entirety.

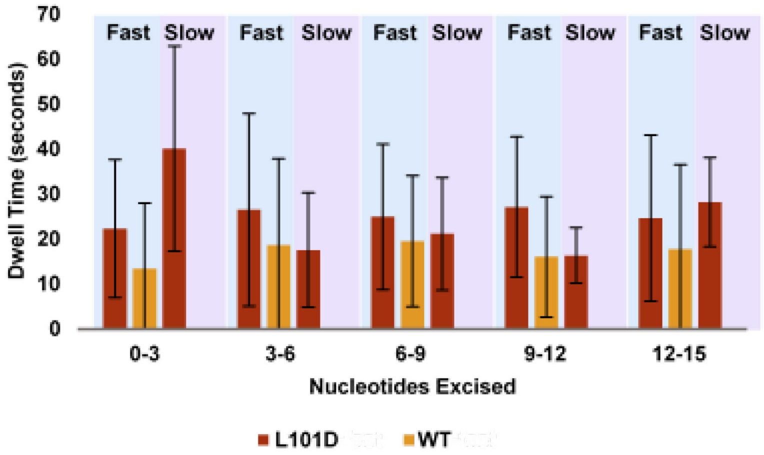


**Figure S9: Dwell time in regards to the overhang length (number of nucleotides already excised).** No correlation between dwell time and the length of the recessed DNA end was observed for either L101D (red) or WT (orange) whether or not the rate subpopulations of slow (purple) or fast (blue) were taken into account (See ANOVA table describiton below).The slow population for WT was too few in number to show here but was included for model fitting for ANOVA. Overhang length was estimated by taking the initial FRET of subsequent transitions and using the linear conversion displayed in Figure S1. This accounts for some error in the estimation of the actual overhang in this data. Note: initial transitions and transitions with a dwell time of zero were not included in this analysis.

**Counts of Dwell Times**

| | WT | | L101D | |
|---|---|---|---|---|
| | **fast** | **slow** | **fast** | **slow** |
| **0-3** | 11 | 2 | 17 | 6 |
| **3-6** | 41 | 0 | 18 | 26 |
| **6-9** | 42 | 2 | 38 | 23 |
| **9-12** | 22 | 1 | 24 | 20 |
| **12-15** | 13 | 1 | 16 | 2 |
| **15-18** | 3 | 0 | 2 | 0 |

| | SS | Df | F value | p value |
|---|---|---|---|---|
| (Intercept) | 109210 | 1 | 414.7401 | <2e-16*** |
| Mutant | 2631 | 1 | 9.9899 | 0.001723** |
| Residuals | 85053 | 323 | | |

**Table S7: ANOVA Table (Type III tests) for Figure S8**

AIC, BIC, and an F-test was used in R to fit the ideal ANOVA model to data shown in Figure S8. The tested variables were overhang length (nucleotides excised prior to interaction), rate subpopulation, and construct. After model fitting, a 1-way ANOVA including only construct (WT- or L101D-MR) was selected indicating that all other main effects are insignificant. Consistently, regardless of other variables, WT dwell times were shorter than L101D dwell times for all overhang lengths. This is consistent with results discussed with Figure 4b and the correlating tables. The length of the overhang has proven to be insignificant according to this statistical analysis. However, the sample sizes for this analysis are small enough that, while the results are a reasonable indicator of the trends of these populations, they may not be conclusive. It should be noted that, due to a low number of WT-slow transitions collected in the study, analysis of the impact of rate subpopulation is limited in this dataset.

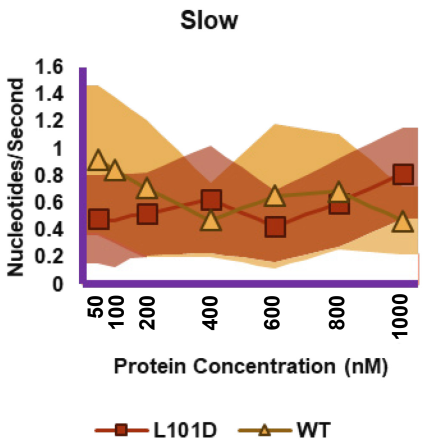


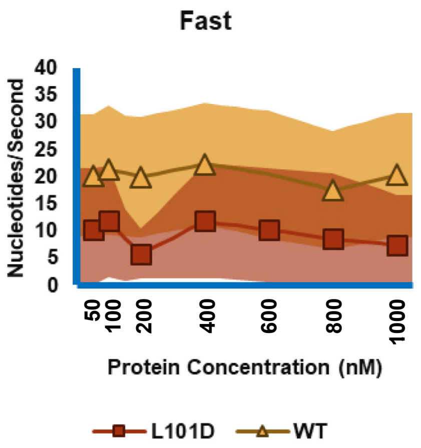


**Figure S10: Rate of Exonuclease Activity by Concentration**
Neither WT-MR (red squares) or L101D-MR (orange triangles) showed a significant correlation between protein concentration and rate even when considering the fast (right, blue) and slow (left, purple) rate subpopulations. The shaded area represents one standard deviation of the of events observed.

**Count of Transitions**

| | WT | | L101D | |
|---|---|---|---|---|
| | **fast** | **slow** | **fast** | **slow** |
| **50** | 119 | 11 | 14 | 43 |
| **100** | 72 | 1 | 49 | 181 |
| **200** | 93 | 6 | 12 | 98 |
| **400** | 110 | 8 | 37 | 221 |
| **600** | 127 | 14 | 7 | 114 |
| **800** | 110 | 28 | 20 | 131 |
| **1000** | 148 | 49 | 35 | 132 |

Table for Figure 5 at different enzyme concentrations

| | **WT (count)** | **L101D (count)** | **WT (%)** | **L101D (%)** |
|---|---|---|---|---|
| **50** | 1 | 0 | 0.008 | 0 |
| **100** | 1 | 1 | 0.014 | 0.004 |
| **200** | 0 | 0 | 0 | 0 |
| **400** | 0 | 1 | 0 | 0.004 |
| **600** | 2 | 0 | 0.014 | 0 |
| **800** | 5 | 0 | 0.036 | 0 |
| **1000** | 0 | 0 | 0 | 0 |

**Supplemental Table S6: Count of Pause Times of Zero**
Fast and slow data were combined. The data does not provide support of increased ratios of zero time transitions with increased concentration. However, a substantially larger quantity of data would be needed to confirm with confidence a significant change in proportions.

Tables with counts:

| | fast | slow |
|---|---|---|
| **L101D** | 490 | 1752 |
| **WT** | 861 | 54 |

Table for Figure S9

| | WT | | L101D | |
|---|---|---|---|---|
| | **fast** | **slow** | **fast** | **slow** |
| **0-3** | 11 | 2 | 17 | 6 |
| **3-6** | 41 | 0 | 18 | 26 |
| **6-9** | 42 | 2 | 38 | 23 |
| **9-12** | 22 | 1 | 24 | 20 |
| **12-15** | 13 | 1 | 16 | 2 |
| **15-18** | 3 | 0 | 2 | 0 |

| | WT | | L101D | |
|---|---|---|---|---|
| | **fast** | **slow** | **fast** | **slow** |
| **Initial** | 724 | 37 | 368 | 1627 |
| **Sub.** | 133 | 6 | 115 | 77 |

Table for Figure 5

| | WT | | L101D | |
|---|---|---|---|---|
| | **Initial** | **Sub.** | **Initial** | **Sub.** |
| **50** | 113 | 16 | 57 | 0 |
| **100** | 65 | 8 | 225 | 5 |
| **200** | 92 | 7 | 109 | 1 |
| **400** | 107 | 11 | 250 | 8 |
| **600** | 120 | 21 | 119 | 2 |
| **800** | 119 | 19 | 151 | 0 |
| **1000** | 184 | 13 | 167 | 0 |

Table for Figure 7, Diffent Enzyme Concentrations

| | WT | | L101D | |
|---|---|---|---|---|
| | **fast** | **slow** | **fast** | **slow** |
| **50** | 119 | 11 | 14 | 43 |
| **100** | 72 | 1 | 49 | 181 |
| **200** | 93 | 6 | 12 | 98 |
| **400** | 110 | 8 | 37 | 221 |
| **600** | 127 | 14 | 7 | 114 |
| **800** | 110 | 28 | 20 | 131 |
| **1000** | 148 | 49 | 35 | 132 |

Table for Figure 6 at different enzyme concentrations
